# Ferrous Iron Accumulation Is a Hallmark and Therapeutic Vulnerability of Therapy-Induced Senescence

**DOI:** 10.64898/2026.05.20.726695

**Authors:** Zhiming Wang, Yuanhui Liu, Hala S. Hassanain, Yaopeng Ding, Shaoyang Zhao, Nancy Azizian, Yinan Gong, Keith S. Chan, Jenny C. Chang, Mark Pegram, Yulin Li

## Abstract

Chemotherapy and radiation reduce tumor burden but leave behind residual cells that survive via therapy-induced senescence (TIS). These cells constitute a latent reservoir fueling recurrence, yet strategies for their selective elimination are lacking. Here, we identify lysosomal ferrous iron accumulation as a conserved hallmark and actionable vulnerability of TIS tumor cells. Across diverse models, senescent tumor cells exhibit marked hypersensitivity to ferroptosis induction. In breast cancer PDX models, sequential ferroptosis induction following chemotherapy significantly delays recurrence, while dual inhibition of GPX4 and FSP1 produces durable, often complete, eradication of residual tumors without overt toxicity. Mechanistically, activation of the TFEB-HO-1 axis in TIS tumor cells drives ferrous iron accumulation, thereby priming cells for ferroptosis. Together, these findings establish ferrous iron accumulation as a defining feature of TIS and position ferroptosis induction as a potent senolytic strategy to eliminate therapy-refractory residual disease.

**Statement of significance:** Senescent tumor cells remaining after treatment can drive cancer recurrence yet remain poorly understood and therapeutically intractable. Here, we identify lysosomal ferrous iron accumulation as a universal hallmark of therapy-induced senescence and demonstrate that ferroptosis induction functions as an effective senolytic strategy. Our findings provide mechanistic and translational support for the “one-two punch” therapeutic paradigm.

## Introduction

Chemotherapy and radiation can substantially reduce tumor burden; however, a subset of tumor cells survives treatment by entering a senescence-like state. This phenomenon, termed <u>T</u>herapy-Induced <u>S</u>enescence (TIS), has been observed *in vitro*, in animal models, and more recently, in patient tumors. TIS cells withstand otherwise lethal doses of genotoxic therapy and persist as dormant, drug-tolerant remnants that can later drive tumor recurrence months or years after apparent clinical remission^1–5^.

TIS shares key features with replicative senescence associated with aging, including enlarged morphology, persistent DNA damage signaling, senescence-associated β-galactosidase (SA-β-gal) activity, and expression of a senescence-associated secretory phenotype (SASP). However, unlike the largely irreversible growth arrest of replicative senescence, TIS represents a reversible and adaptive state^3–5^. Tumor cells that undergo TIS can eventually escape from growth arrest, regain proliferative capacity, and drive tumor recurrence. Thus, selectively targeting TIS tumor cells with senolytic strategies represents a promising approach to prevent recurrence and limit the emergence of more aggressive, therapy-resistant disease. Despite this potential, clinical translation has been limited, in part due to an incomplete understanding of the molecular mechanisms governing TIS and its associated vulnerabilities.

To address this gap, we previously established an *in vitro* model that recapitulates the senescent phenotype observed in post-chemotherapy human tumors^6^. A chemical library screen encompassing approximately 250 non-genotoxic agents revealed distinct vulnerabilities in TIS cells, particularly within pathways regulating ferroptosis, apoptosis, and necroptosis^6^. Building on these findings, we conducted a secondary, focused screen of cell death modulators in an independent TIS model^4^, and validated ferroptosis as a selective and robust mechanism for eliminating senescent tumor cells. Using patient-derived xenograft (PDX) models of triple-negative breast cancer (TNBC), we demonstrate that ferroptosis induction effectively eradicates post-chemotherapy residual tumors and significantly delays recurrence. Notably, concurrent inhibition of GPX4 and FSP1, two parallel ferroptosis defense systems, produces durable, and in some cases complete, tumor regression without discernible toxicity. Mechanistically, we identify activation of TFEB-HO-1 axis and consequent lysosomal ferrous iron accumulation as key determinants of ferroptosis hypersensitivity in TIS cells. This adaptive stress response to cancer therapy paradoxically primes senescent cells for iron-dependent ferroptotic death.

Collectively, our findings identify ferrous iron accumulation as a defining hallmark of TIS tumor cells and establish ferroptosis induction as a potent senolytic strategy to eliminate residual disease and prevent tumor recurrence.

## Results

### Ferroptosis emerges as a selective vulnerability of therapy-induced senescent tumor cells

To identify targetable vulnerabilities in therapy-induced senescence (TIS), we first leveraged our previously established *in vitro* model in which chemotherapy combined with mTOR inhibition induces a robust and reversible senescent state across multiple cancer types^6^. These TIS cells display canonical features of senescence, including enlarged morphology, elevated senescence-associated β-galactosidase (SA-β-gal) activity, induction of p21 and γH2AX, lamin B1 loss, and a typical senescence-associated secretory phenotype (SASP)^6^. Importantly, these TIS models were reversible, as senescent tumor cells resumed proliferation within approximately two weeks of drug withdrawal^6^.

An initial chemical library screen with 250 compounds targeting diverse cellular processes identified multiple pathways selectively lethal to TIS, but not control cells, including G2/M checkpoint regulation, autophagy/lysosome function, and cell death pathways^6^. To more directly interrogate cell death dependencies, we performed a secondary focused screen of approximately 80 compounds targeting apoptosis, necroptosis, ferroptosis, pyroptosis, and cuproptosis in an independent TIS model derived from MDA-MB-231 cells (**Fig. 1a-d**)^4^. Strikingly, TIS cells exhibited pronounced hypersensitivity to multiple ferroptosis inducers, including RSL3, ML162, ML210, and JKE-1716, suggesting that ferroptosis represents a dominant and previously underappreciated vulnerability of the senescent state (**Fig. 1d**, **Supplementary Table 1**).

**Fig 1.**
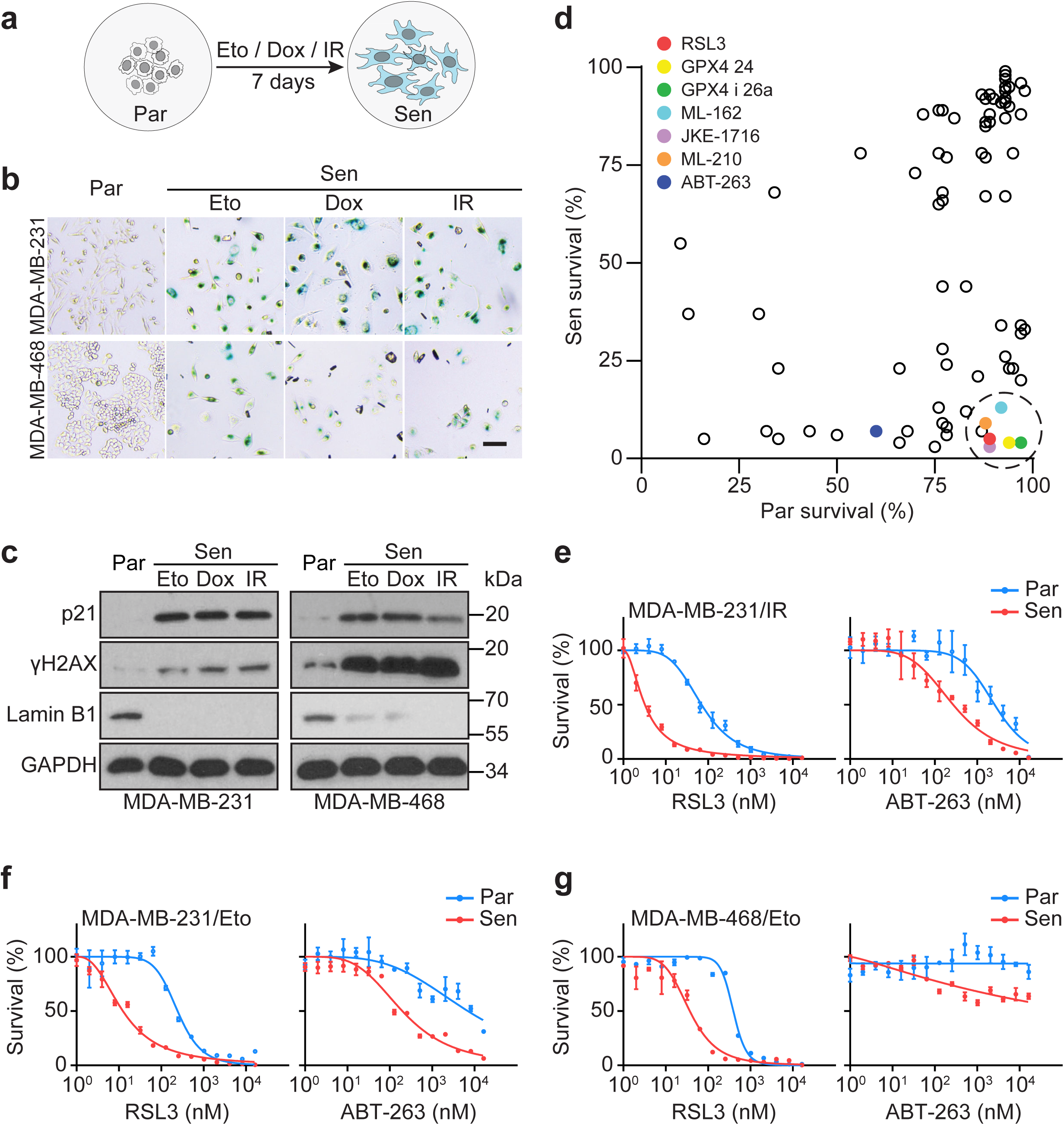
A chemical library screen identifies ferroptosis inducers as effective senolytics. **a.** Schematic overview of senescence induction by chemotherapy or irradiation. Par, parental; Sen, senescent; Eto, 1.25 μM etoposide; Dox, 75 nM doxorubicin; IR, 10 Gy X-ray irradiation. **b.** SA-β-gal staining of MDA-MB-231 and MDA-MB-468 TIS cells generated by chemotherapy (1.25 μM etoposide and 75 nM doxorubicin for MDA-MB-231; 0.75 μM etoposide and 25 nM doxorubicin for MDA-MB-468) or X-ray irradiation (10 Gy). Scale bar, 100 μm. **c.** Western blot analysis of p21, γH2AX, and Lamin B1 levels in MDA-MB-231 and MDA-MB-468 TIS cells. GAPDH served as a loading control. **d.** Chemical library screen in MDA-MB-231 TIS cells. The dashed circle indicates ferroptosis-inducing hits. Percent survival of parental and senescent cells following treatment with various cell death modulators was normalized to untreated controls. **e-g.** Sensitivity of TIS cells derived from X-ray irradiation (**e**) or etoposide (**f-g**) to RSL3 and ABT-263 after 24 hours of treatment. Data represent the mean of four technical replicates; error bars indicate standard deviation.

### Ferroptosis induction broadly and selectively eliminates TIS tumor cells

We next sought to validate ferroptosis as a senolytic strategy across diverse contexts. Treatment with the GPX4 inhibitor RSL3 efficiently eliminated TIS cells induced by chemotherapy or X-ray irradiation in multiple TNBC models, while largely sparing proliferating controls (**Fig. 1e-g**). Notably, RSL3 displayed substantially greater potency than the canonical senolytic agent ABT-263 at low nanomolar concentrations^4^.

Mechanistically, RSL3 treatment triggered a marked increase in lipid peroxidation in TIS cells, as measured by BODIPY 581/591 C11^7,8^, consistent with ferroptotic cell death. This effect was specifically suppressed by ferroptosis inhibitors, but not by inhibitors of apoptosis or necroptosis, confirming ferroptosis as the dominant mode of cell death (**Supplementary Fig. 1a-c**).

Importantly, ferroptosis sensitivity was not restricted to a single treatment modality or cancer type. TIS cells induced by multiple chemotherapeutic agents, including doxorubicin, irinotecan, 5-FU, gemcitabine, and cisplatin exhibited similar hypersensitivity to RSL3 (**Supplementary Fig. 2**). This vulnerability extended across a broad range of cancer models, including ER+ and HER2+ breast cancers, as well as pancreatic, lung, colorectal, prostate, and melanoma cell lines. Across these models, RSL3 consistently outperformed ABT-263 in both potency and selectivity (**Supplementary Fig. 3-4**). Together, these findings establish ferroptosis induction as a robust and broadly applicable senolytic strategy for TIS tumor cells.

### Ferroptosis induction suppresses tumor recurrence in vivo

We next evaluated whether ferroptosis induction could eliminate senescent residual tumors *in vivo*. Using treatment-naïve TNBC PDX models, we induced tumor regression with a standard doxorubicin-cyclophosphamide (AC) regimen, resulting in a population of residual tumors that displayed hallmark features of senescence, including increased p21, γH2AX, and lamin B1 loss (**Fig. 2a-e**)^9^.

**Fig 2.**
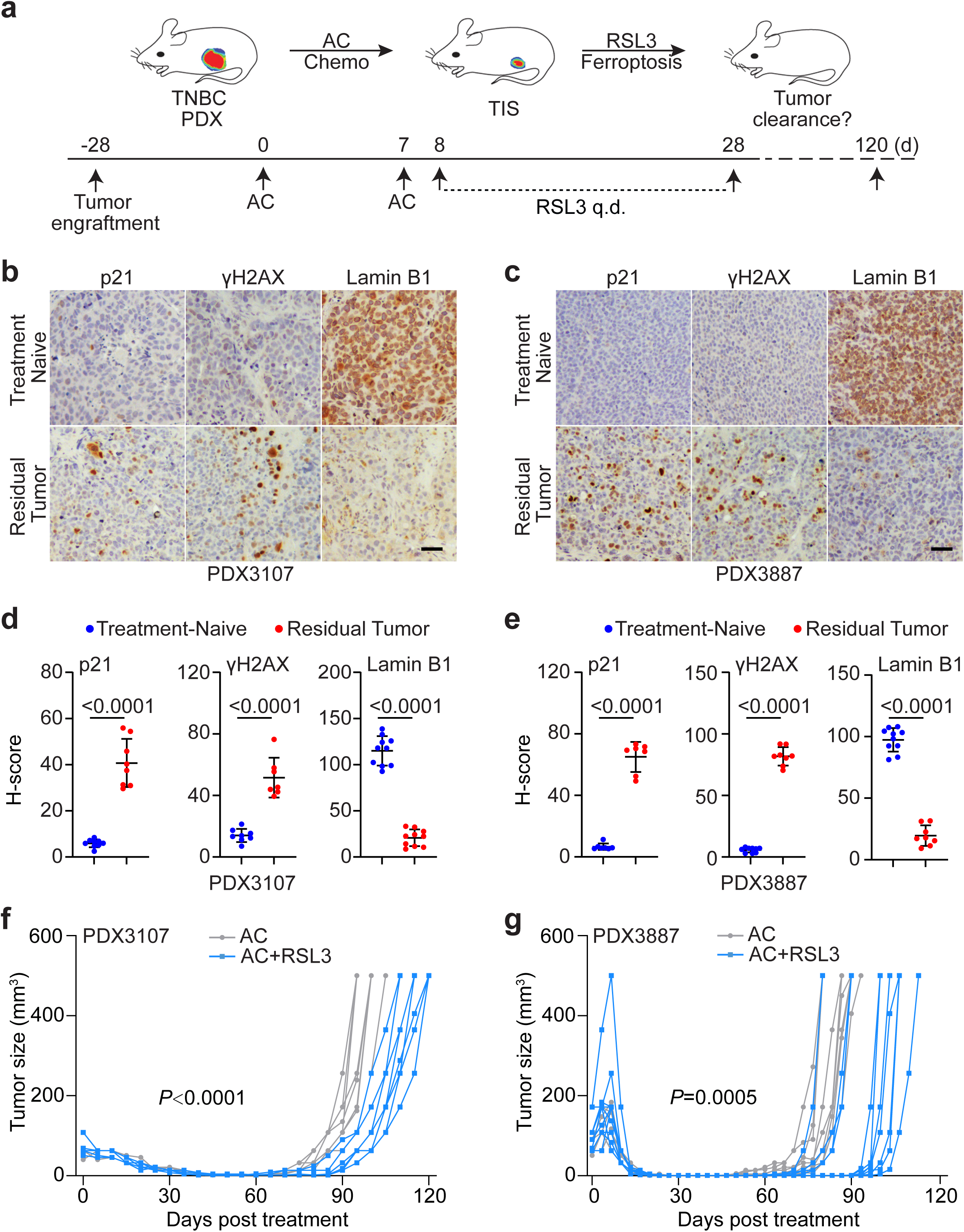
Ferroptosis induction by RSL3 significantly delays tumor recurrence. **a.** Schematic illustration of sequential ferroptosis induction in post-chemotherapy residual tumors. Residual tumors were established following two to three cycles of AC (doxorubicin, 0.5 mg/kg; cyclophosphamide 50 mg/kg, i.p.). Daily RSL3 treatment (three weeks) was initiated after the final AC dose. **b-e.** Immunohistochemical staining of p21, γH2AX, and Lamin B1 in residual tumors from PDX3107 and PDX3887 collected 21 days after chemotherapy. H-score quantification was performed using ImageJ. Scale bar, 100 μm. **f-g.** Sequential RSL3 treatment following chemotherapy delays tumor recurrence in PDX3107 and PDX3887. P values represent differences in time-to-recurrence between AC and AC+RSL3 groups (Student’s t-test). Source data are provided in Supplementary Table 2a.

Sequential treatment with RSL3 following chemotherapy significantly delayed tumor recurrence compared to chemotherapy alone, extending recurrence-free survival with no detectable changes in body weight or overt toxicity. These findings demonstrate that ferroptosis induction can function as an effective senolytic strategy *in vivo* to suppress post-therapy tumor relapse (**Fig. 2f-g, Supplementary Fig. 5a-c, Supplementary Table 2a**). However, despite this benefit, tumors eventually recurred in all cases following RSL3 monotherapy, suggesting the existence of intrinsic resistance mechanisms limiting ferroptosis efficacy.

### Dual inhibition of GPX4 and FSP1 eradicates senescent tumors and induces durable remission

Given the incomplete responses to GPX4 inhibition alone, we hypothesized that parallel ferroptosis defense pathways may limit therapeutic efficacy. FSP1 is known to function independently of GPX4 to suppress lipid peroxidation and confer resistance to ferroptosis^10–12^. Suppression of FSP1 using chemical inhibitors (iFSP1, icFSP1, FSEN1) synergizes with GPX4 blockade to promote ferroptosis^13,14^.

Consistent with this model, combined inhibition of GPX4 and FSP1 using RSL3 and iFSP1 produced strong synergistic cytotoxicity in vitro, selectively eradicating senescent tumor cells more effectively than either agent alone. Notably, this synergy was substantially more pronounced in TIS cells than in proliferating controls, underscoring the selective ferroptosis vulnerability of the senescence state (**Supplementary Fig. 6a-b**).

We next evaluated this combination *in vivo* using a panel of TNBC PDX TIS models. Across all models, the RSL3/iFSP1 combination markedly outperformed RSL3 monotherapy, resulting in prolonged recurrence-free survival and significantly extended overall survival. Importantly, in several PDXs, including PDX3887, PDX0089, PDX0140, and PDX0060, dual GPX4/FSP1 inhibition achieved complete and durable tumor eradication, with a majority of treated animals remaining tumor-free for up to 11 months following cessation of therapy (**Fig. 3a-f, Supplementary Fig. 7a, Supplementary Table 2b**). Residual tumors treated with RSL3 or RSL3/iFSP1 exhibited significantly increased lipid peroxidation based on staining of 4-HNE (4-Hydroxynonenal) and MDA (Malondialdehyde) (**Fig. 3g-h**)^15^. Notably, the combination did not cause obvious toxicity, as assessed by animal appearance/behavior, body weight, and histopathology (**Supplementary Fig. 7b-c**).

**Fig 3.**
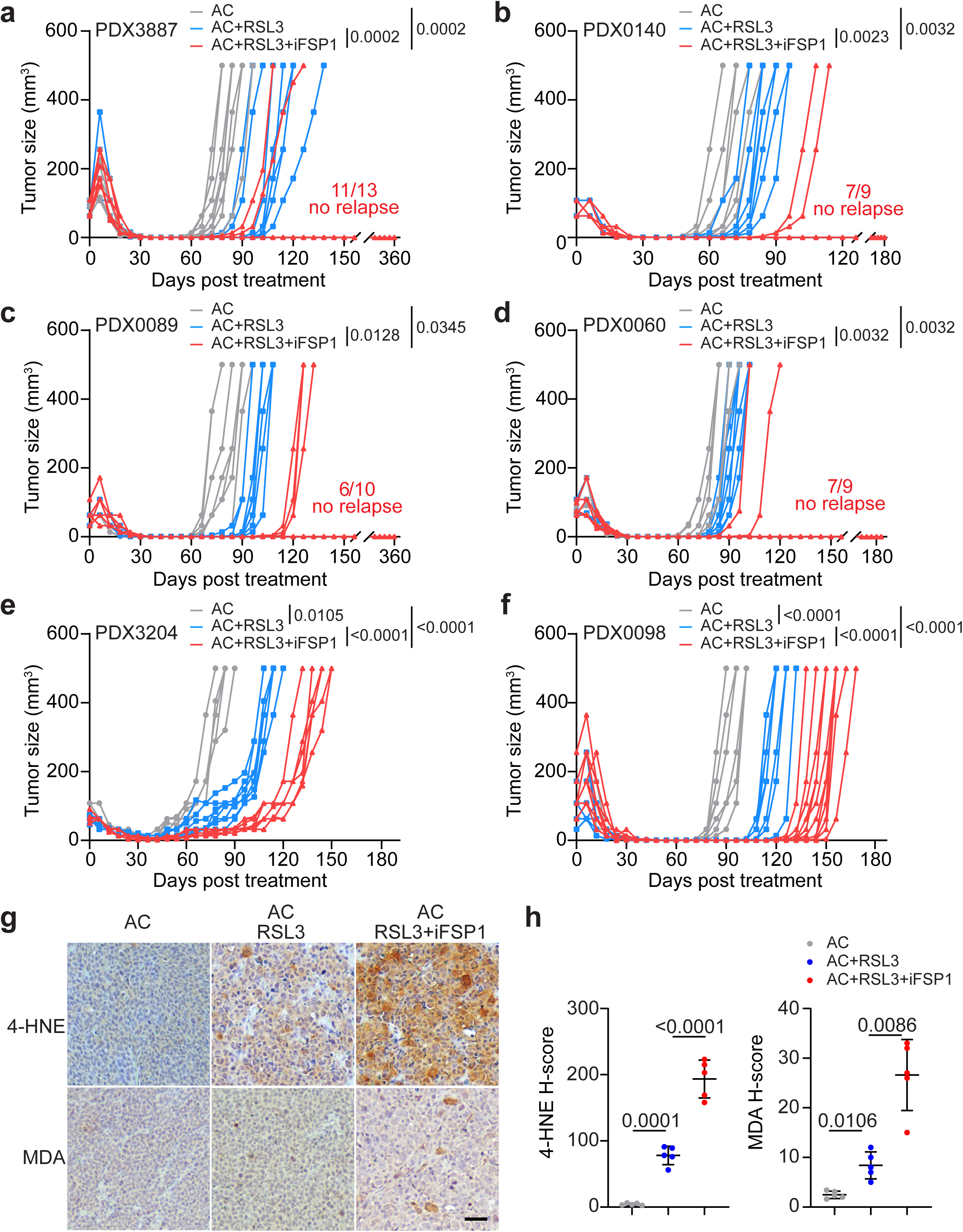
Co-suppression of GPX4 and FSP1 is highly synergistic *in vivo* in eradicating TIS tumors. **a-f.** *In vivo* therapeutic efficacy of combined RSL3 and iFSP1 across a panel of TNBC PDX residual tumor models. The number of cures in each treatment group is shown. Source data are provided in **Supplementary Table 2b**. For PDX3204 and PDX0098, time-to-recurrence was analyzed using one-way ANOVA. For PDX3887, PDX0140, PDX0089, and PDX0060, recurrence frequencies were compared using Fisher’s exact test. **g-h**. IHC staining of lipid peroxidation markers, 4-HNE and MDA, in tumors treated with chemotherapy alone, chemotherapy plus RSL3, or chemotherapy plus RSL3/iFSP1. Tumors were collected after nine days of RSL3 or RSL3/iFSP1 treatments. P values (one-way ANOVA) comparing H-scores are shown. Scale bar, 100 μm.

To assess broader therapeutic applicability, we extended ferroptosis induction to residual tumors in additional cancer types, including colorectal cancer (PDX0170, irinotecan), HER2+ breast cancer (PDX0261, T-Dxd), and acute myeloid leukemia (luciferase-labeled U937, cytarabine). Ferroptosis induction, via combined RSL3/iFSP1 treatment in the solid tumor models, and RSL3 alone in U937, substantially reduced residual tumor burden and delayed recurrence (**Supplementary Fig. 8a-f**).

Taken together, these results demonstrate that ferroptosis induction, particularly when combined with inhibition of compensatory defense pathways, can achieve durable remission with the potential for curative outcomes.

### Lysosomal ferrous iron accumulation is a defining hallmark of TIS tumor cells

To understand the basis for ferroptosis hypersensitivity in TIS, we examined intracellular iron metabolism. Ferrous iron is a key driver of lipid peroxidation through the Fenton reaction and is therefore a central determinant of ferroptosis sensitivity. Using the FerroOrange probe^16^, we observed a consistent 3-5-fold increase in intracellular ferrous iron levels in TIS cells induced by chemotherapy or radiation. This increase was conserved across multiple cancer types and treatment conditions (**Fig. 4a, Supplementary Figs. 9-10**).

**Fig 4.**
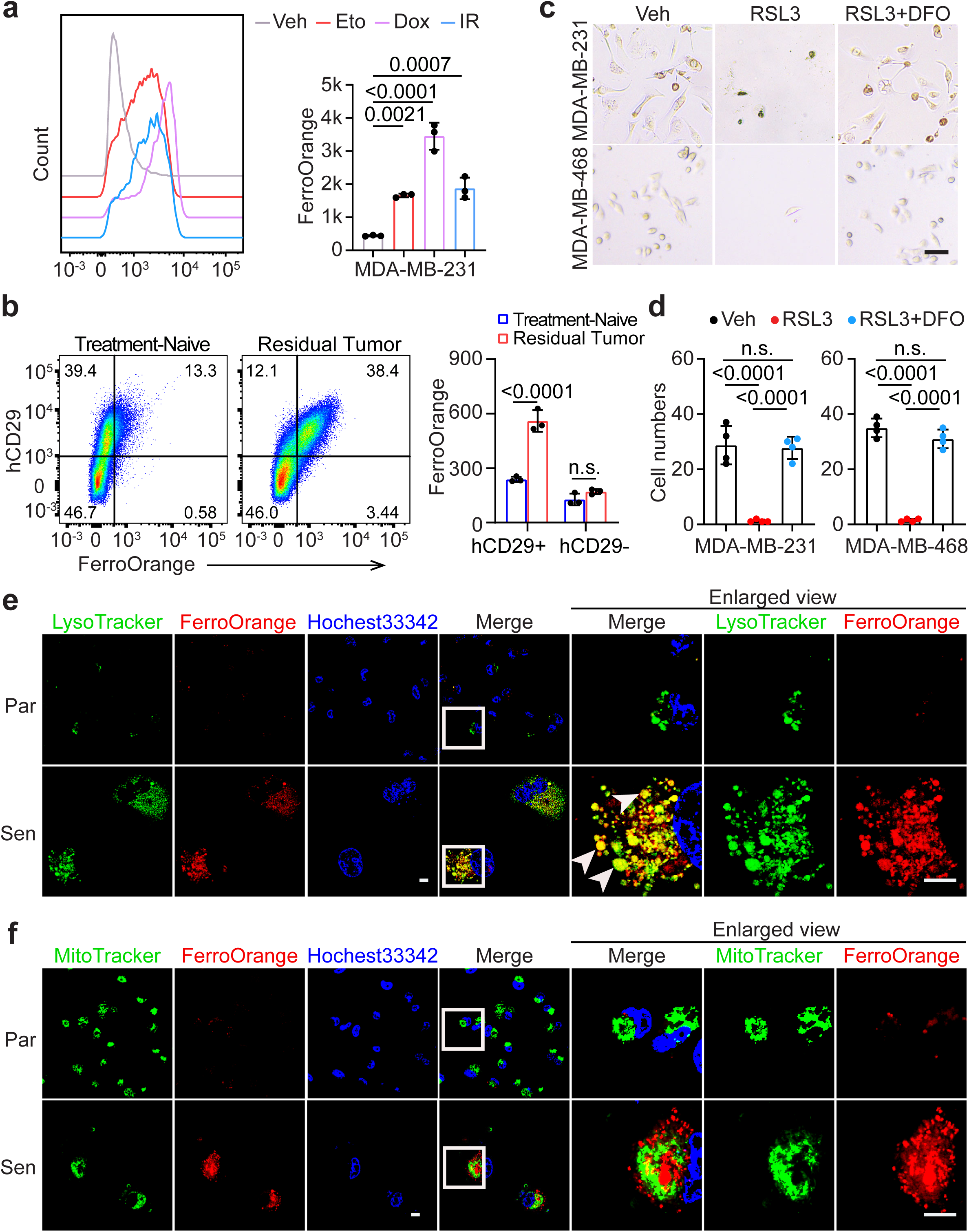
Ferrous iron accumulation within lysosomes confers ferroptosis hypersensitivity to TIS cells. **a.** Quantification of intracellular ferrous iron levels in parental and TIS MDA-MB-231 cells by flow cytometry following FerroOrange staining. P values (one-way ANOVA) comparing each condition with parental cells are shown. **b.** *In vivo* ferrous iron levels in tumor *versus* stromal cells from treatment-naïve and AC-treated PDX3887 models. Tumors from three treatment-naïve and AC-treated mice were dissected and dissociated into single-cell suspensions, and human CD29 staining was used to gate tumor cells. Residual tumors were collected 21 days after chemotherapy. P values were determined by one-way ANOVA. n.s., not significant. **c-d.** The iron chelator deferoxamine (DFO) prevents RSL3-induced ferroptosis. MDA-MB-231 and MDA-MB-468 TIS cells were treated with RSL3 (100 nM) with or without DFO (10-100 µM) for 24 hours. The data shown corresponds to DFO at 10 µM. P values (one-way ANOVA) are shown. n.s., not significant. **e-f.** Confocal imaging of intracellular ferrous iron in parental and TIS MDA-MB-231 cells. Cells were stained with FerroOrange (ferrous iron), Hoechst 33342 (nuclei), LysoTracker (lysosomes), and MitoTracker (mitochondria). Arrowheads indicate regions of overlapping FerroOrange and LysoTracker signals. Scale bar, 20 μm.

Importantly, we further quantified ferrous iron in single-cell suspensions of treatment-naïve and post-chemotherapy residual tumors from the PDX3887 model. By gating on human CD29 (hCD29, integrin β1)^17^, we detected an approximately three-fold increase in ferrous iron levels specifically within residual tumor cells, but not in the hCD29-negative stromal compartment, indicating that iron accumulation is a tumor cell-intrinsic feature of the TIS state (**Fig. 4b**).

Functionally, iron chelation with deferoxamine or deferasirox abrogated ferroptosis induction, demonstrating that elevated ferrous iron is required for this vulnerability (**Fig. 4c-d, Supplementary Fig. 11**). Notably, confocal imaging and flow cytometric analysis revealed expanded lysosomal compartments in TIS cells, with ferrous iron localized predominantly to lysosomes rather than mitochondria (**Fig. 4e-f**).

To assess specificity, we compared TIS with other non-proliferative and drug-tolerant states. Quiescent tumor cells induced by serum starvation or mTOR inhibition exhibited reduced ferrous iron levels despite minimal changes in lysosomal content (**Supplementary Fig. 12a-b**). While oncogene-targeted therapies can induce drug-tolerant persisters (DTP), they generally do not robustly induce senescence. Notably, multiple DTP models generated by oncogene-targeted therapies failed to exhibit increased ferrous iron levels (**Supplementary Fig. 12c-f**). These findings are particularly notable given prior reports that DTPs can remain sensitive to ferroptosis^18,19^. Our results therefore demonstrate that iron loading is a TIS-specific metabolic feature rather than a general property of drug tolerance. Collectively, these data establish lysosomal ferrous iron accumulation as a defining and distinguishing hallmark of TIS tumor cells.

### The TFEB-HO-1 axis drives iron accumulation and ferroptosis sensitivity in TIS

We next sought to identify the mechanisms underlying ferrous iron accumulation in TIS. Among known iron-releasing pathways, we focused on ferritinophagy and heme degradation. Ferritinophagy, mediated by NCOA4, degrades ferritin to liberate iron^20^. Heme oxygenases catalyze the breakdown of intracellular heme into ferrous iron, carbon monoxide, and biliverdin^21^. Of the two heme oxygenases, HO-1 (encoded by *HMOX1*) is stress-inducible, whereas HO-2 (encoded by *HMOX2*) is constitutively expressed^21^. Notably, HO-1 is induced by chemotherapy and radiation across cancer types, and has been implicated in cellular adaptation and therapy resistance^21–25^. Western blot analysis showed robust and consistent induction of HO-1 in TIS cells, whereas the levels of ferritinophagy components, ferroptosis suppressors, and HO-2 remained largely unchanged across multiple TIS models induced by chemotherapy and X-ray irradiation (**Fig. 5a, Supplementary Fig. 13a-b**). HO-1 induction was validated *in vivo* in post-chemotherapy residual tumors from PDX3107 and PDX3887 models (**Fig. 5b-c**). Collectively, these data suggest that HO-1 is a potential driver of ferrous iron accumulation in TIS cells.

**Fig 5.**
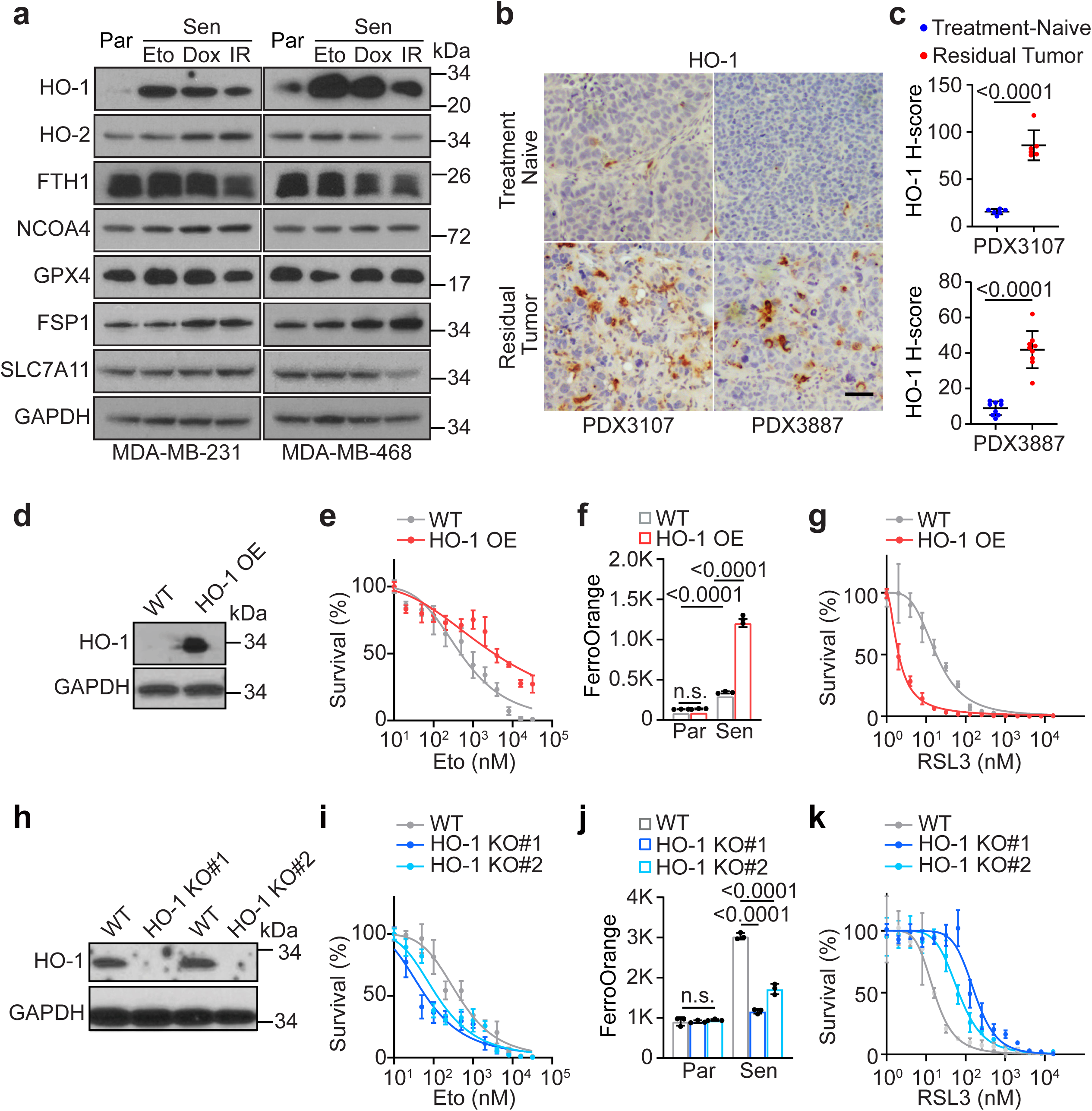
HO-1 activation promotes ferrous iron accumulation and ferroptosis sensitivity. **a.** Western blot analysis of heme oxygenases, ferritinophagy regulators, and ferroptosis suppressor proteins in MDA-MB-231 and MDA-MB-468 TIS cells. **b-c.** Immunohistochemical detection of HO-1 expression in residual tumor cells from PDX3107 and PDX3887. P values (Student’s t-test) comparing H-scores are shown. Scale bar, 100 μm. **d.** Lentiviral HO-1 overexpression (OE) in MDA-MB-231 cells confirmed by western blot analysis. GAPDH served as a loading control. **e.** HO-1 overexpression induces chemoresistance. Cells were treated with increasing concentrations of etoposide for seven days, and the surviving fraction was quantified by CellTiter-Glo assay. **f-g.** HO-1 overexpression promotes ferrous iron accumulation (**f**) and increases ferroptosis sensitivity (**g**) in TIS cells. Cells were treated with etoposide for seven days, followed by measurement of ferrous iron levels using FerroOrange staining and flow cytometric analysis. Ferroptosis sensitivity in senescent cells was assessed after one day of RSL3 treatment. P values (one-way ANOVA) are shown above the bar graphs. **h.** CRISPR knockout of *HMOX1* gene in MDA-MB-231 cells confirmed by western blot analysis. Two knockout clones (KO#1, KO#2) were identified. GAPDH served as a loading control. **i.** HO-1 knockout clones exhibit increased chemosensitivity. Cells were treated with increasing concentrations of etoposide for seven days, and the surviving fraction was quantified using CellTiter-Glo assay. **j-k.** CRISPR knockout of *HMOX1* gene blocks ferrous iron accumulation following senescence induction (**j**) and dampens ferroptosis sensitivity (**k**). Wild-type and knockout clones were treated with etoposide for seven days, followed by measurement of ferrous iron levels using FerroOrange staining and flow cytometric analysis. Ferroptosis sensitivity in senescent wild-type and knockout cells was assessed after one day of RSL3 treatment. P values (one-way ANOVA) are shown above the bar graphs.

To validate the functional role of HO-1, we genetically manipulated its encoding gene, *HMOX1*, in MDA-MB-231 cells. *HMOX1* overexpression increased ferrous iron levels and enhanced ferroptosis sensitivity, while conferring resistance to chemotherapy. Conversely, genetic ablation of *HMOX1* prevented iron accumulation and suppressed ferroptosis induction in TIS cells (**Fig. 5d-k**). Together, these findings demonstrate that HO-1 activation is a major functional driver of iron-dependent ferroptosis hypersensitivity in TIS tumor cells.

Upstream of HO-1, we identified activation of TFEB, a master regulator of lysosomal biogenesis and stress responses^26–29^. In three residual tumors models, including two TNBC PDXs^9^, and one lung cancer cell line-derived xenograft model (CDX-H441)^30^, we observed pronounced nuclear TFEB staining in residual tumor cells but not treatment-naïve tumor cells, indicating activation of the TFEB pathway following chemotherapy (**Fig. 6a-b**).

**Fig 6.**
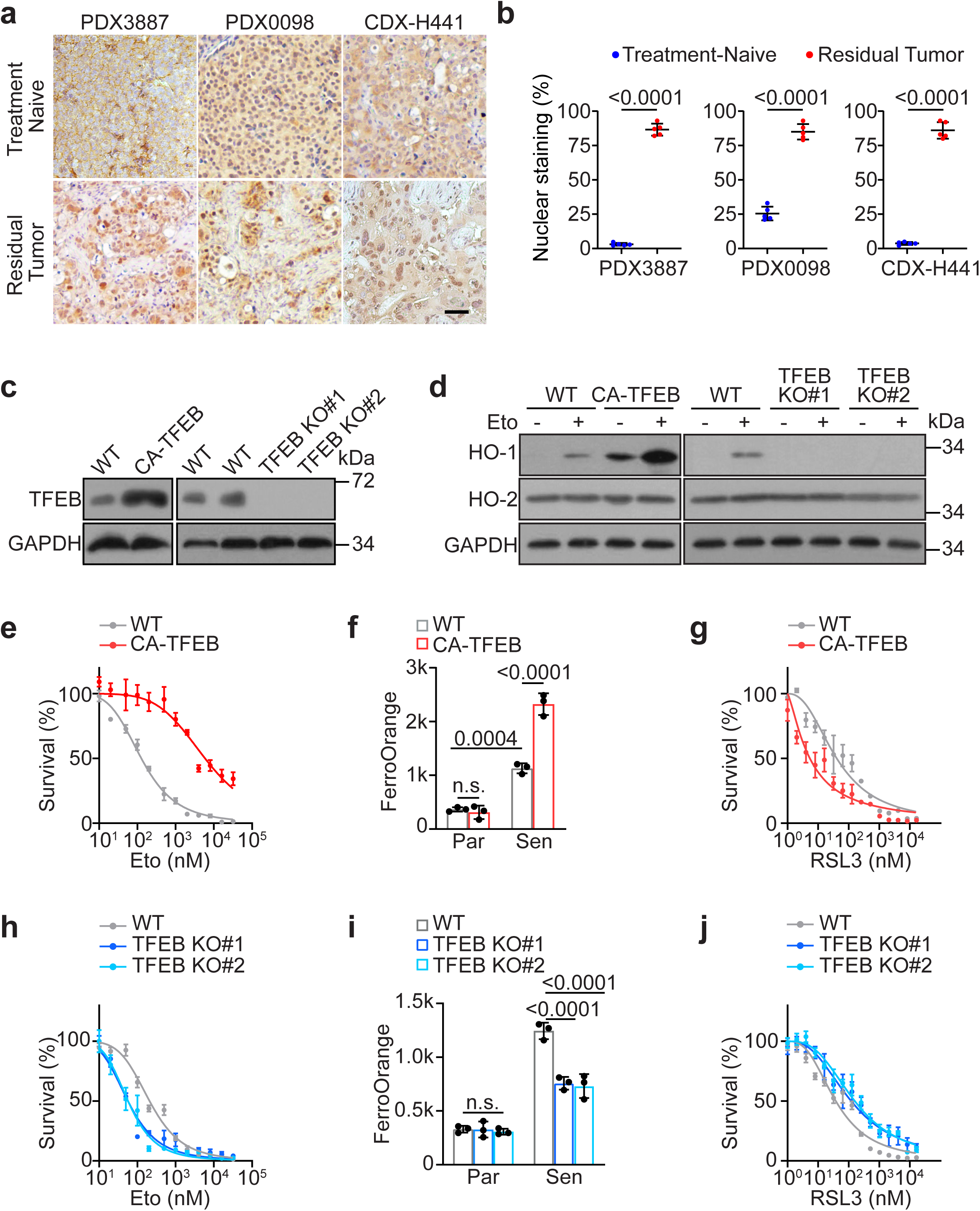
TFEB regulates HO-1 expression and modulates ferrous iron accumulation and ferroptosis sensitivity. **a-b.** Immunohistochemical detection of TFEB protein expression in treatment-naïve versus post-chemotherapy residual tumor cells derived from PDX3887, PDX0098, and CDX-H441. PDX3887 and PDX0098 were treated AC as described previously. CDX-H441 was treated with a combination of paclitaxel (20 mg/kg, i.p., every two days for 10 days) and cisplatin (5 mg/kg, i.p., weekly for two weeks). P values (Student’s t-test) comparing H-scores are shown. Scale bar, 100 μm. **c.** Western blot analysis of MDA-MB-231 cells following lentiviral expression of constitutively active TFEB (CA-TFEB) or CRISPR-mediated *TFEB* knockout. GAPDH served as a loading control. **d.** Changes in HO-1 and HO-2 protein expression levels following genetic manipulation of TFEB activity. GAPDH served as a loading control. **e.** CA-TFEB expression induces chemoresistance. Cells were treated with increasing concentrations of etoposide for seven days, and the surviving fraction was quantified by CellTiter-Glo assay. **f-g.** CA-TFEB expression promotes ferrous iron accumulation (**f**) and ferroptosis sensitivity (**g**) in TIS cells. Cells were treated with etoposide for seven days, followed by measurement of ferrous iron levels using FerroOrange staining and flow cytometric analysis. Ferroptosis sensitivity in senescent cells was assessed after 24 hours of RSL3 treatment. P values (one-way ANOVA) are shown above the bar graphs. **h.** *TFEB* knockout clones exhibit increased chemosensitivity. Cells were treated with increasing concentrations of etoposide for seven days, and the surviving fraction was quantified by CellTiter-Glo assay. **i-j.** CRISPR-mediated knockout of *TFEB* gene decreases ferrous iron accumulation following senescence induction (**i**) and dampens ferroptosis sensitivity (**j**). Wild-type (WT) and knockout (KO) clones were treated with etoposide for seven days; ferrous iron levels were measured via FerroOrange staining and flow cytometry. Ferroptosis sensitivity was assessed after 24 hours of RSL3 treatment. P values (one-way ANOVA) are shown above the bar graphs.

To determine whether HO-1 and downstream iron accumulation are directly regulated by TFEB, we utilized lentiviral expression of a constitutively active TFEB (CA-TFEB)^29^ and CRISPR-mediated knockout of TFEB (**Fig. 6c**). Constitutive TFEB activation promoted HO-1 induction, whereas TFEB knockout blocked chemotherapy-induced HO-1 expression; notably, HO-2 levels remained unaffected by these genetic manipulations (**Fig. 6d**). Furthermore, TFEB activation conferred chemoresistance, increased ferrous iron accumulation, and enhanced ferroptosis sensitivity. Conversely, TFEB ablation promoted chemosensitivity, reduced iron accumulation, and dampened ferroptosis induction (**Fig. 6e-j**). Taken together, these data indicate that activation of the TFEB-HO-1 axis is a major mechanism underlying ferrous iron accumulation and ferroptosis sensitivity in TIS tumor cells.

## Discussion

We demonstrate that senescent tumor cells across diverse cancer types and treatment modalities consistently exhibit marked lysosomal ferrous iron accumulation. This metabolic shift directly underlies their heightened sensitivity to ferroptosis. In multiple PDX models, the sequential administration of ferroptosis inducers following chemotherapy significantly delayed tumor recurrence without observable systemic toxicity. Notably, the concurrent inhibition of GPX4 and FSP1, two complementary and partially redundant defense pathways, more effectively eradicated residual senescent cells, achieving complete and durable remission in some instances. Together, these findings identify ferrous iron accumulation as a conserved hallmark of TIS and establish ferroptosis induction as a potent senolytic strategy. Furthermore, our results provide mechanistic and translational support for the “one-two punch” paradigm, in which senescence induction is followed by selective elimination of senescent tumor cells to prevent recurrence^1,2^.

Mechanistically, our data implicate the TFEB-HO-1-ferrous iron axis as a central mechanistic link between drug tolerance and ferroptosis hypersensitivity in many TIS settings. TFEB-HO-1 activation after cancer therapy elevates the lysosomal ferrous iron pool, thereby priming senescent tumor cells for ferroptotic cell death. While we focused on GPX4 and FSP1, emerging strategies that directly activate lysosomal iron pools, such as the lysosome-targeting agent fentomycin-1, may further potentiate ferroptosis in TIS tumors^31^. Future investigations into lysosomal iron trafficking and redox regulation may uncover additional upstream regulators and therapeutic targets for selectively eliminating senescent tumor cells.

While the ferroptosis inducers used in this study (RSL3 and iFSP1) are tool compounds that lack optimized pharmacokinetic properties for clinical use, their favorable tolerability and strong efficacy in multiple PDX models provide compelling proof-of-concept evidence that systemic ferroptosis induction can selectively target senescent residual disease *in vivo*. The absence of overt toxicity indicates a wide therapeutic window driven by the unique iron biology of TIS cells compared with normal tissues. These results may motivate the development of next-generation ferroptosis inducers with improved pharmacokinetic properties for clinical translation.

A key conceptual advance of this study is the distinction between therapy-induced senescence and other drug-tolerant states at the level of iron metabolism. Although prior studies have demonstrated that drug-tolerant persister (DTP) cells following targeted therapy can be susceptible to ferroptosis^18,19^, we find that DTPs do not exhibit increased intracellular ferrous iron levels; indeed, others have reported reduced iron abundance in these cells^18^. These observations indicate that ferroptosis sensitivity can arise independently of iron loading. In contrast, TIS cells undergo robust TFEB-HO-1-driven iron accumulation, creating a distinct metabolic state that amplifies ferroptotic vulnerability. Thus, iron loading, rather than ferroptosis sensitivity *per se*, represents the defining and functionally important hallmark unique to TIS.

Finally, the consistency of ferrous iron accumulation across diverse TIS models suggests that iron dysregulation is a core and conserved feature of the senescent state. Given that elevated ferrous iron levels have also been reported during organismal aging in mammals and *C. elegans*^32–35^, our work raises the possibility that iron accumulation may represent a shared metabolic feature contributing to senescence across biological contexts. Targeting this metabolic vulnerability may therefore have broad implications not only for cancer therapy, but also for aging and age-associated diseases.

## Methods

### Cell lines and chemical reagents

MDA-MB-231, MDA-MB-468, T47D, MCF7, AU565, BT474, A375, AsPC-1, MIA PaCa-2, HCT116, H441, U937, SW480, PC3, LNCaP and A549 were from ATCC. All cells were cultured in DMEM supplemented with 10% fetal bovine serum and 1% penicillin/streptomycin in a 5% CO2 humidified incubator. Doxorubicin and cyclophosphamide were obtained from our institutional pharmacy. RSL3 was purchased from Selleckchem. iFSP1, KT-253, alisertib, and all targeted therapeutic agents were purchased from MedChemExpress. All chemicals for the small-molecule library screening were purchased from Cayman Chemical.

### In vitro models of therapy-induced senescence

We followed previously published protocols from Dr. René Bernards’s laboratory to establish *in vitro* TIS models using specific chemotherapeutic agents and concentrations^4^. For radiation-induced senescence, tumor cells were exposed to 10 Gy X-ray irradiation. Senescence was confirmed by characteristic morphological changes, western blot analysis of senescence markers (p21, γH2AX, and Lamin B1), and SA-β-gal staining. Tumor cell quiescence was induced by either serum starvation (0.1% fetal bovine serum) or mTOR inhibition (100 nM Torin1) for seven days.

### *In vitro* analysis of response to RSL3 and ABT-263

After seven days of treatment, the cells reached approximately 70-90% confluency and exhibited a senescent phenotype. For analysis of the response to RSL3 and ABT-263, parental and senescent cells collected from T300 flasks were reseeded into 96-well plates. The following day, cells were treated with either RSL3 or ABT-263 across a concentration range of 1 nM to 16 µM using 1:2 serial dilutions for 24 hours. Cell viability was assessed using CellTiter-Glo (Promega) in technical quadruplicates.

### SA-*β*-gal staining

After seven days of drug treatment, cells were stained using the Senescence Cells Histochemical Staining Kit (Sigma, CS0030) according to the manufacturer’s instructions. Nine images per well were acquired, and positively stained cells were manually counted.

### Chemical library screen using cell death modulators

Approximately 80 chemical inhibitors targeting major cell-death pathways were dissolved in DMSO or PBS. Inhibitors were generally tested in a 1:3 dilution series from 10 µM to 100 nM, although some were tested at concentrations as low as 30 nM. Parental and senescent MDA-MB-231 cells were treated with individual inhibitors for five days. Concentrations that produced significant viability changes in either parental or senescent cells were recorded. Viability data (average of three to four fields) were plotted at the concentrations showing the greatest differences. If both parental and senescent cells were killed, experiments were repeated using lower concentrations. For inhibitors characterized in previous studies, we consulted original publications and adjusted the concentration ranges accordingly.

### Western blot analysis

Cells were lysed on ice using CelLytic MT reagent (Sigma) supplemented with protease and phosphatase inhibitors. Lysates were subjected to SDS-PAGE, transferred to PVDF membranes, and immunoblotted. The following antibodies were used: p21 (#2947, CST, 1:1000), Lamin B1 (#12586, CST, 1:1000), γH2AX (#2577, CST, 1:2000), HO-1 (#43966, CST, 1:1000), HO-2 (#32790, CST, 1:1000), TFEB (#37785, CST, 1:1000), GPX4 (#52455, CST, 1:1000), FSP1 (#24972, CST, 1:1000), SLC7A11 (#98051, CST, 1:1000), NCOA4 (#66849, CST, 1:1000), FTH1 (#3998, CST, 1:1000), and GAPDH (#10494-1-ap, ProteinTech, 1:10000).

### Flow cytometry analysis

Lipid peroxidation was assessed using the fluorescent probe C11-BODIPY 581/591 (#95978, CST) following established protocols. After exposure to RSL3 for one to two hours, parental and senescent cells were stained with 2 µM C11-BODIPY 581/591 in serum-containing medium for 20 minutes at 37 °C, washed, and analyzed. Intracellular ferrous iron was measured using the FerroOrange probe (F374, Dojindo, 1 µM) according to the manufacturer’s instructions. Samples were analyzed on a BD FACS LSRII cytometer (BD Biosciences), collecting at least 10,000 events per sample. Standard gating strategies were applied to exclude debris and doublets. For simultaneous quantification of lysosomal mass and intracellular ferrous irons, cells were treated with LysoTracker Green DND-26 (#8783, CST, 50 nM) and FerroOrange (F374, Dojindo, 1 µM). All data were analyzed using FlowJo v10.8.1 (Tree Star).

### Confocal imaging analysis

For visualization of intracellular ferrous iron, cells were stained with FerroOrange (F374, Dojindo, 1 µM), Hoechst 33342 (R37605, Thermo Scientific, 1 µg/mL), LysoTracker Green DND-26 (#8783, CST, 50 nM), and MitoTracker Green FM (#9074, CST, 50 nM) for 20 minutes at 37 °C. Images were acquired on an Olympus FluoView FV3000 confocal laser scanning microscope with appropriate filter sets. ImageJ software was used for image analysis.

### In vivo residual tumor studies

For therapeutic studies, PDX models were obtained from the Jackson Laboratory (PDX0098: TM00098; PDX0089: TM00089; PDX0170: TM00170), Baylor College of Medicine (PDX3107: BCM3107; PDX3887: BCM3887; PDX3204: BCM3204), and NCI Patient-Derived Models Repository (PDX0140: 994819 140 R; PDX0060: 755484-060-R; PDX0261: K68440 261 T). Tumor size was measured with calipers when the diameter exceeded 4 mm, or by palpation for tumors smaller than 4 mm. Tumor volume was calculated as a*b^2^/2, where a and b are tumor length and width. For the luciferase-labeled U937 (U937-luc) acute myeloid leukemia model, bioluminescence imaging (BLI) was performed using an IVIS Lumina III system and analyzed with Living Image software (v4.2). RSL3 was dissolved in DMSO to prepare a 500 mg/mL stock solution, which was subsequently diluted in a vehicle containing 5% DMSO and 95% corn oil for intraperitoneal injection at 30 mg/kg. iFSP1 was dissolved in DMSO to generate a 160 mg/mL stock solution and further diluted in a vehicle composed of 5% DMSO, 40% PEG300, 5% Tween-80, and 50% saline for oral gavage at 10 mg/kg. Time-to-recurrence was defined as the days by which the tumor regrows to pre-treatment volumes. All mice were housed under standard conditions (ambient temperature, 60% humidity, 12-hour light/dark cycle) with ad libitum access to food and water. Mice were euthanized when tumor diameter reached 1.5 cm. All procedures were approved by the Houston Methodist Research Institute IACUC and complied with institutional and national guidelines.

### Isolation of single cells from tumors for flow cytometric analysis

Treatment-naïve tumors and post-AC residual tumors were minced and digested into single-cell suspensions using tumor dissociation kits and processed on the gentleMACS dissociator (Miltenyi Biotec). Suspensions were filtered through 70-µm strainers, centrifuged at 300 × g for 5 minutes, and resuspended for staining with FerroOrange (1 µM for 20 minutes) and an Alexa Fluor 488 anti-human CD29 antibody (BioLegend, #303016, 1:100 for 20 minutes).

### Immunohistochemical analysis of tumor sections

Formalin-fixed, paraffin-embedded tissue sections were deparaffinized, and endogenous peroxidase activity was quenched with 3% hydrogen peroxide for 10 minutes. Sections were blocked with 5% normal serum for 30 minutes, followed by overnight incubation at 4 °C with primary antibodies against p21, γH2AX, Lamin B1, TFEB, HO-1 (all from CST, 1:300), MDA (JAI-MMD-030N, AdipoGen, 1:100), and 4-HNE (ab46545, Abcam, 1:200). After washing with PBS, sections were incubated with an HRP-conjugated secondary antibody for 30 minutes at room temperature. Signal detection was performed using a DAB substrate kit (#8059, CST), and nuclei were briefly counterstained with hematoxylin. Slides were then dehydrated, cleared, and mounted with coverslips. ImageJ was used to quantify staining intensity and percentage of cells stained using the H-score method.

### Genetic manipulation of HO-1 and TFEB expression

For HO-1 expression, the HO-1 coding sequence from pCX-HO1-2A-EGFP (Addgene #74672) was cloned into the lentiviral vector pLV-EF1a-IRES-Puro (Addgene #85132). To express the constitutively active CA-TFEB (S142A, S211A), the coding sequence was cloned into the lentiviral vector pLV-EF1a-IRES-Blast (Addgene #85133). Lentiviral particles were produced in 293FT cells and used to infect MDA-MB-231 cells via spin infection.

For construction of sgRNA-expressing vectors, annealed DNA oligonucleotides were cloned into either the lentiGuide-Puro vector (Addgene #52963) for *HMOX1* or PX458 (Addgene #48138) for *TFEB*. The target sgRNA sequences are listed in **Supplementary Table 3**. Lentiviral particles encoding the *HMOX1* sgRNAs were generated in 293FT cells and delivered into MDA-MB-231 cells stably expressing Cas9 (Addgene #52962) by spin infection. Three days after puromycin selection, cells were seeded into 96-well plates for single-cell cloning to isolate HO-1 knockout lines. In contrast, *TFEB* sgRNAs were delivered by electroporation; GFP-positive cells were then isolated by fluorescence-activated cell sorting into 96-well plates as single cells. The loss of HO-1 and TFEB protein expression was validated by western blot analysis.

### Statistical analysis

Animal survival was assessed using Kaplan–Meier survival curves and compared using the log-rank (Mantel-Cox) test. For quantitative comparisons between groups, statistical significance was determined using an unpaired two-tailed Student’s t-test or one-way ANOVA with a post hoc test, as appropriate. Fisher’s exact test was used to evaluate differences in tumor recurrence rates. All statistical analyses were performed using GraphPad Prism v8.4.2.

## Data availability

All source data are included within the paper. All other materials are available on request from the corresponding author.

## Acknowledgements

Y.Li. was supported by the Cancer Prevention and Research Institute of Texas (CPRIT; RP200472), the NIH (R01CA288884), and the American Cancer Society (RSG-24-1322678-01-TBE). H.H. was supported by a Burroughs Wellcome Fund Physician Scientist Institutional Award to the Texas A&M University Academy of Physician Scientists. J.C.C. was supported by the NIH (R01CA284315 and U01CA268813).

## Author contributions

Z.W. performed most of the *in vitro* and *in vivo* experiments and analyzed the data. Y.Liu identified ferroptosis hypersensitivity and supported data analyses. H.H. contributed to the identification of HO-1-ferrous iron axis. S.Z. assisted with the PDX studies. N.A. contributed to the FerroOrange assays and PDX modeling. J.C.C. and M.P. advised the study. Y.Li. conceived and supervised the project and wrote the manuscript.

## Supplementary Figure Legends

**Supplementary Fig 1. Senolytic activity of RSL3 occurs through ferroptosis.**

**a.** Detection of lipid peroxidation using BODIPY 581/591 C11 in parental and TIS cells. Parental and TIS MDA-MB-231 and MDA-MB-468 cells were treated with RSL3 (100 nM, 1-2 hours) and stained with BODIPY 581/591 C11 (2 µM, 20 minutes) for flow cytometric analysis. P values from one-way ANOVA are shown above the bar graphs.

**b-c**. Effects of cell-death pathway inhibitors on the senolytic activity of RSL3. TIS MDA-MB-231 cells were treated with RSL3 (100 nM) in the presence or absence of inhibitors of ferroptosis (ferrostatin-1, 100 nM; liproxstatin-1, 100 nM), necroptosis (necrostatin-1s, 10 µM), and apoptosis (Z-VAD-FMK, 10 µM). Representative images are shown in **b**, and quantification is shown in **c**. Live cells from at least three 20x fields were counted. P values from one-way ANOVA are shown.

**Supplementary Fig 2. RSL3 and ABT-263 sensitivity in MDA-MB-231 TIS models induced by diverse chemotherapies.**

MDA-MB-231 cells were treated with 5-FU (**a**, 50 µM), gemcitabine (**b**, 100 nM), irinotecan (**c**, 5 µM), cisplatin (**d**, 5 µM), oxaliplatin (**e**, 10 µM), or doxorubicin (**f**, 150 nM) to generate the TIS models. Sensitivity to RSL3 and ABT-263 was assessed as described above.

**Supplementary Fig 3. RSL3 and ABT-263 sensitivity in ER+ and HER2+ breast cancer TIS models.**

**a-b**. ER+ breast cancer (MCF7, T47D) and HER2+ breast cancer (AU565, BT474) TIS models induced by etoposide, doxorubicin, or X-ray irradiation. Treatment conditions: MCF7 (2.5 µM Eto; 100 nM Dox; 10 Gy IR), T47D (0.5 µM Eto; 75 nM Dox; 10 Gy IR), AU565 (2.5 µM Eto; 50 nM Dox; 10 Gy IR), BT474 (3 µM Eto; 50 nM Dox; 10 Gy IR).

**c-d**. Western blot analysis of p21, γH2AX, and Lamin B1 in ER+ and HER2+ TIS cells. GAPDH served as a loading control.

**e-h**. Sensitivity of chemotherapy-induced TIS cells to RSL3 and ABT-263 after 24 hours of treatment. Data represent the mean of four technical replicates; error bars denote standard deviation.

**Supplementary Fig 4. RSL3 and ABT-263 sensitivity in multiple solid tumor TIS models.**

A panel of solid tumor cell lines was treated with the indicated chemotherapies for seven days to induce TIS. Treatments: (**a**) A375, 3 µM etoposide; (**b**) AsPC-1, 50 µM gemcitabine; (**c**) HCT116, 3 µM etoposide; (**d**) SW480, 3 µM etoposide; (**e**) PC3, 100 nM doxorubicin; (**f**) LNCaP, 4 µM etoposide; (**g**) A549, 1.25 µM etoposide; and (**h**) PC9, 4 µM etoposide. Sensitivity to RSL3 and ABT-263 was assessed after 24 hours of treatment.

**Supplementary Fig 5. Overall survival, time-to-recurrence, and body weight changes in animals treated with vehicle or RSL3 alone.**

**a.** Survival analysis of PDX3107 and PDX3887. P values from log-rank tests are shown.

**b.** Time-to-recurrence in AC versus AC+RSL3 groups. Data are presented as mean +/- standard deviation. P values from Student’s t-tests are shown.

**c.** Body-weight changes in animals treated with AC versus AC+RSL3. n.s., not significant.

**Supplementary Fig 6. Co-suppression of GPX4 and FSP1 using RSL3 and iFSP1 is highly synergistic in killing senescent TNBC cells *in vitro*.**

**a-b**. *In vitro* dose-matrix combination of RSL3 and iFSP1 in parental and senescent MDA-MB-231 (**a**) and MDA-MB-468 cells (**b**). Numbers within the matrix indicate the percentage of cell death (cell killing) following 24 hours of treatment.

**Supplementary Fig 7. Overall survival, body weight, and histological changes in animals treated with vehicle, RSL3 alone, or RSL3/iFSP1.**

**a.** Survival analysis for each PDX model. P values from log-rank tests are shown.

**b.** Body weight changes in each PDX cohort across the three treatment conditions. n.s., not significant (one-way ANOVA).

**c.** H&E staining of major organs from untreated (UT) animals or those treated with AC, AC plus RSL3, or AC plus RSL3/iFSP1. Tumors were collected after nine days of RSL3 or RSL3/iFSP1 treatment. Scale bar, 100 μm.

**Supplementary Fig 8. Ferroptosis induction in residual tumor models of colorectal cancer, HER2+ breast cancer, and acute myeloid leukemia.**

**a-b**. *In vivo* efficacy of combined RSL3 and iFSP1 in colorectal cancer (PDX0170) and HER2+ breast cancer (PDX0261). When tumors reached approximately 5 mm in diameter, PDX0170 was treated with irinotecan (45 mg/kg, i.p., twice per week for eight weeks), and PDX0261 was treated with T-DXd (10 mg/kg, intravenous injection, every three weeks for two doses) to generate residual tumors prior to ferroptosis induction with RSL3/iFSP1. P values from one-way ANOVA (time-to-recurrence) are shown.

**c-d**. Survival analyses corresponding to the treatment groups in panels **a-b**. P values from log-rank tests are shown.

**e.** Schematic of sequential ferroptosis induction in a luciferase-labeled U937 (U937-luc) residual tumor model. U937 leukemia-bearing mice were treated with cytarabine (Ara-C; 60 mg/kg, i.p., daily for five days) to induce tumor regression. RSL3 (30 mg/kg, daily for seven days) was initiated after the third Ara-C dose.

**f.** Tumor burden in the U937 leukemia model quantified by bioluminescence imaging (BLI). P value from Student’s t-test is shown.

**Supplementary Fig 9. Ferrous iron levels in ER+ and HER2+ breast cancer TIS models.**

**a.** Gating strategy for flow cytometric measurement of ferrous iron in parental and TIS cells.

**b.** Ferrous iron levels in ER+ and HER2+ breast cancer TIS models. Samples were analyzed in biological triplicates; P values from one-way ANOVA are shown.

**Supplementary Fig 10. Ferrous iron levels in TIS models derived from multiple solid tumor types treated with various chemotherapeutic agents.**

**a-h**. Ferrous iron measurements in the solid tumor TIS models described in **Supplementary Fig 4**. Samples were analyzed in biological triplicates; P values from Student’s t-test are shown.

**Supplementary Fig 11. Iron chelation blocks RSL3-induced ferroptosis in a dose-dependent manner.**

**a-b**. Representative images and quantification of live MDA-MB-231 TIS cells following RSL3 treatment (100 nM for 24 hours) in the presence or absence of increasing concentrations of iron chelators, including deferoxamine (DFO) (**a**) and deferasirox (DFX) (**b**). P values from one-way ANOVA are shown.

**Supplementary Fig 12. Intracellular ferrous iron levels and lysosomal mass and in quiescence and oncogene-targeted therapy models.**

**a-b**. Intracellular ferrous iron levels (FerroOrange) and lysosomal mass (LysoTracker) in quiescent MCF7 and MDA-MB-231 tumor cells induced by serum starvation (0.1%) and mTOR inhibition (100 nM Torin1) for seven days. P values from one-way ANOVA are shown. n.s., not significant.

**c-f**. Ferrous iron levels and lysosomal mass quantified by flow cytometry in MIA PaCa-2 pancreatic cancer cells treated with the KRAS^G12C^ inhibitor adagrasib (Ada, 2-4 µM), A375 melanoma cells treated with the BRAF^V600E^ inhibitor vemurafenib (Vem, 5-10 µM), HER2+ BT474 breast cancer cells treated with the HER2 inhibitor lapatinib (Lap, 2-5 µM), and PC9 lung cancer cells treated with the EGFR inhibitor gefitinib (Gef, 0.2-0.4 µM). All treatments were for seven days. P values from one-way ANOVA are shown. n.s., not significant.

**Supplementary Fig 13. Western blot analysis of heme oxygenases and ferroptosis regulators in additional genotoxic breast cancer TIS models.**

**a-b.** Western blot analysis of heme oxygenases, ferritinophagy regulators, and ferroptosis suppressor proteins in ER+ and HER2+ breast cancer cells following chemotherapy or X-ray irradiation.

## Supplementary Tables

**Supplementary Table 1.** List of cell death modulators used in the chemical library screening.

**Supplementary Table 2.** *In vivo* therapeutic outcome for animals treated with ferroptosis inducers.

**2a.** Days-to-recurrence in PDX3107 and PDX3887 treated with RSL3 alone.

**2b.** Days-to-recurrence in six TNBC PDX models treated with RSL3 or RSL3/iFSP1.

**Supplementary Table 3.** sgRNA sequences used to generate *HMOX1* and *TFEB* CRISPR knockouts.

