## Supplementary Figures 1-13 for "Ferrous Iron Accumulation Is a Hallmark and Therapeutic Vulnerability of Therapy-Induced Senescence"

### Supplementary Fig 1

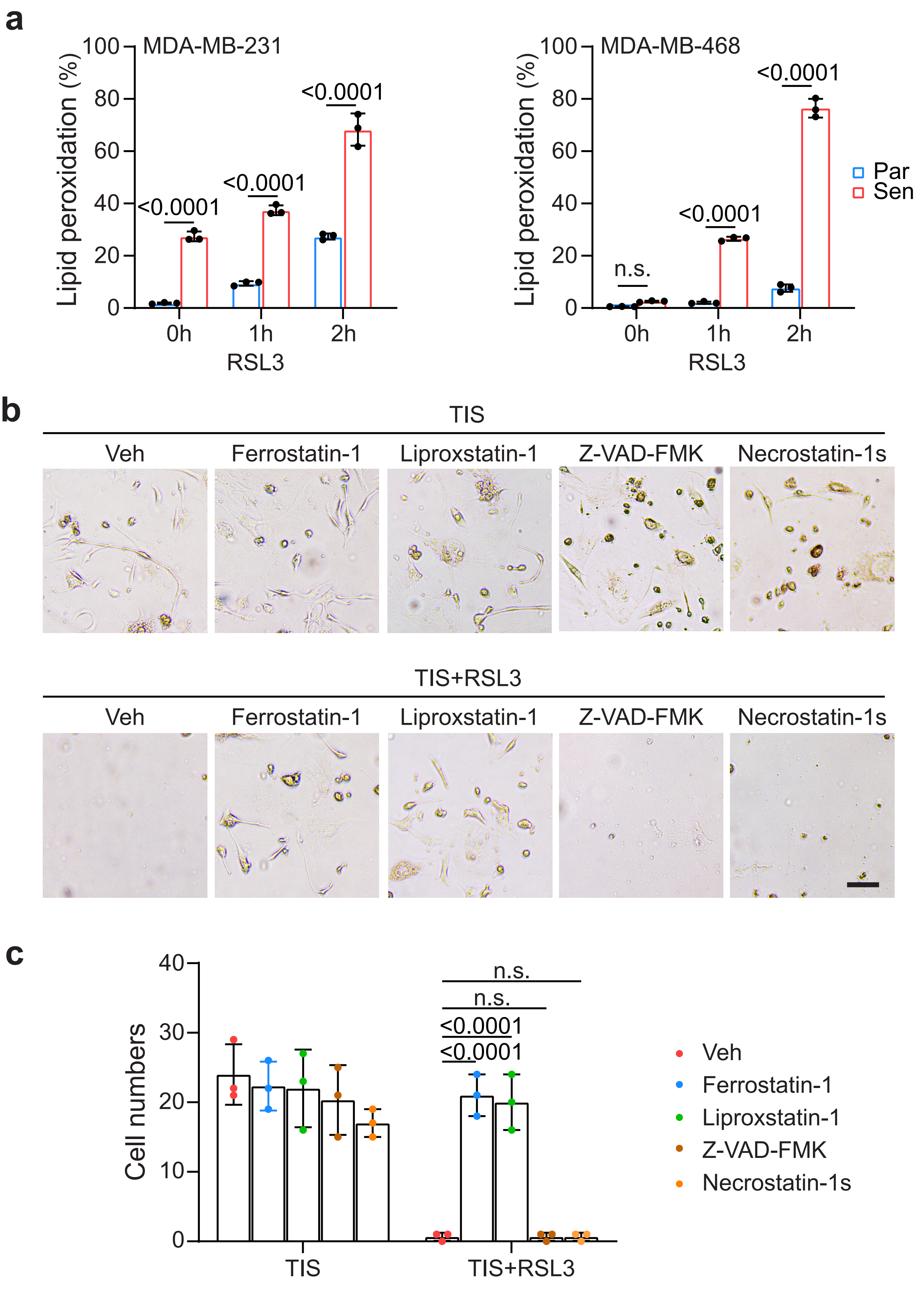

### Supplementary Fig 2

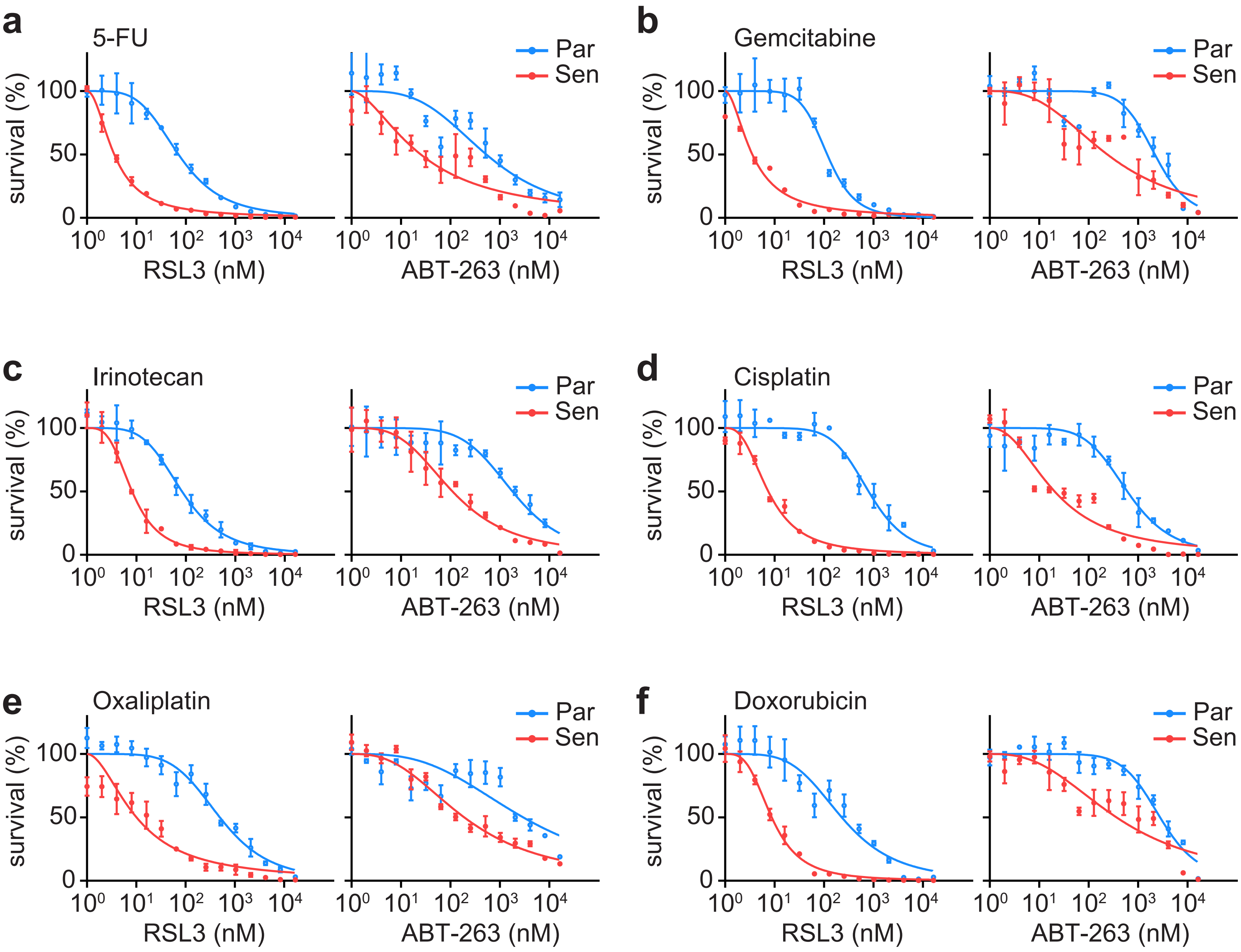

### Supplementary Fig 3

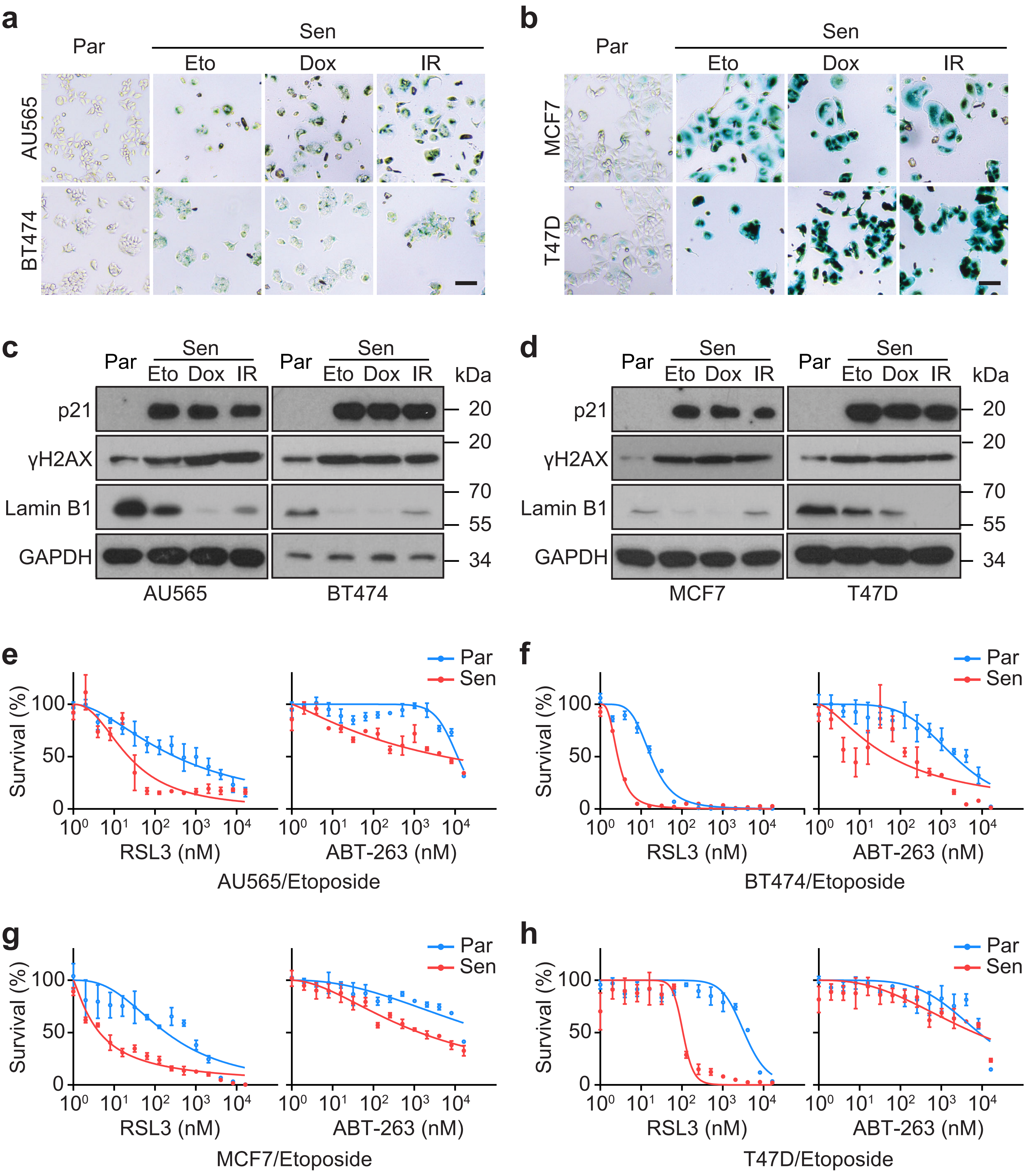

### Supplementary Fig 4

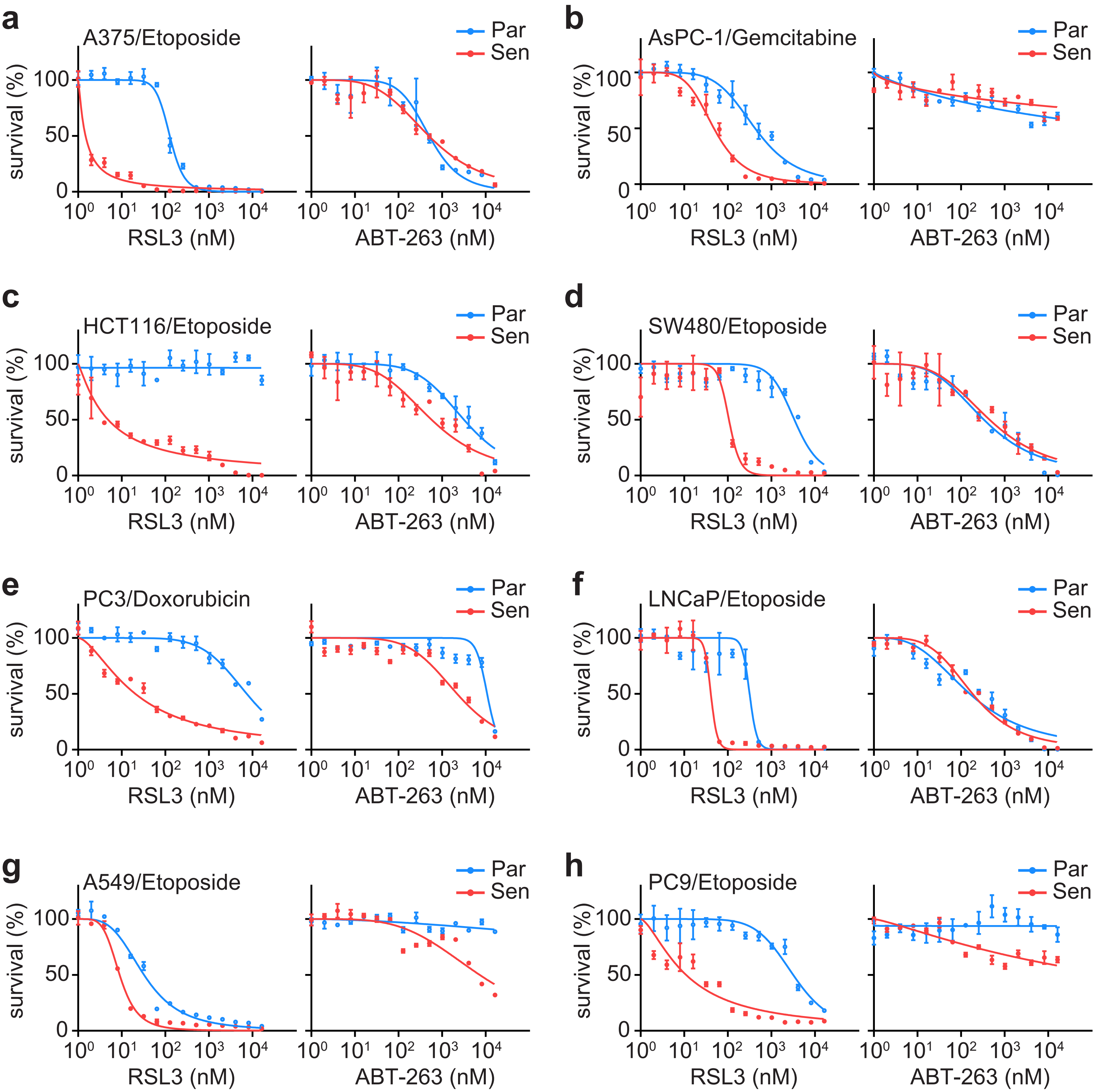

### Supplementary Fig 5

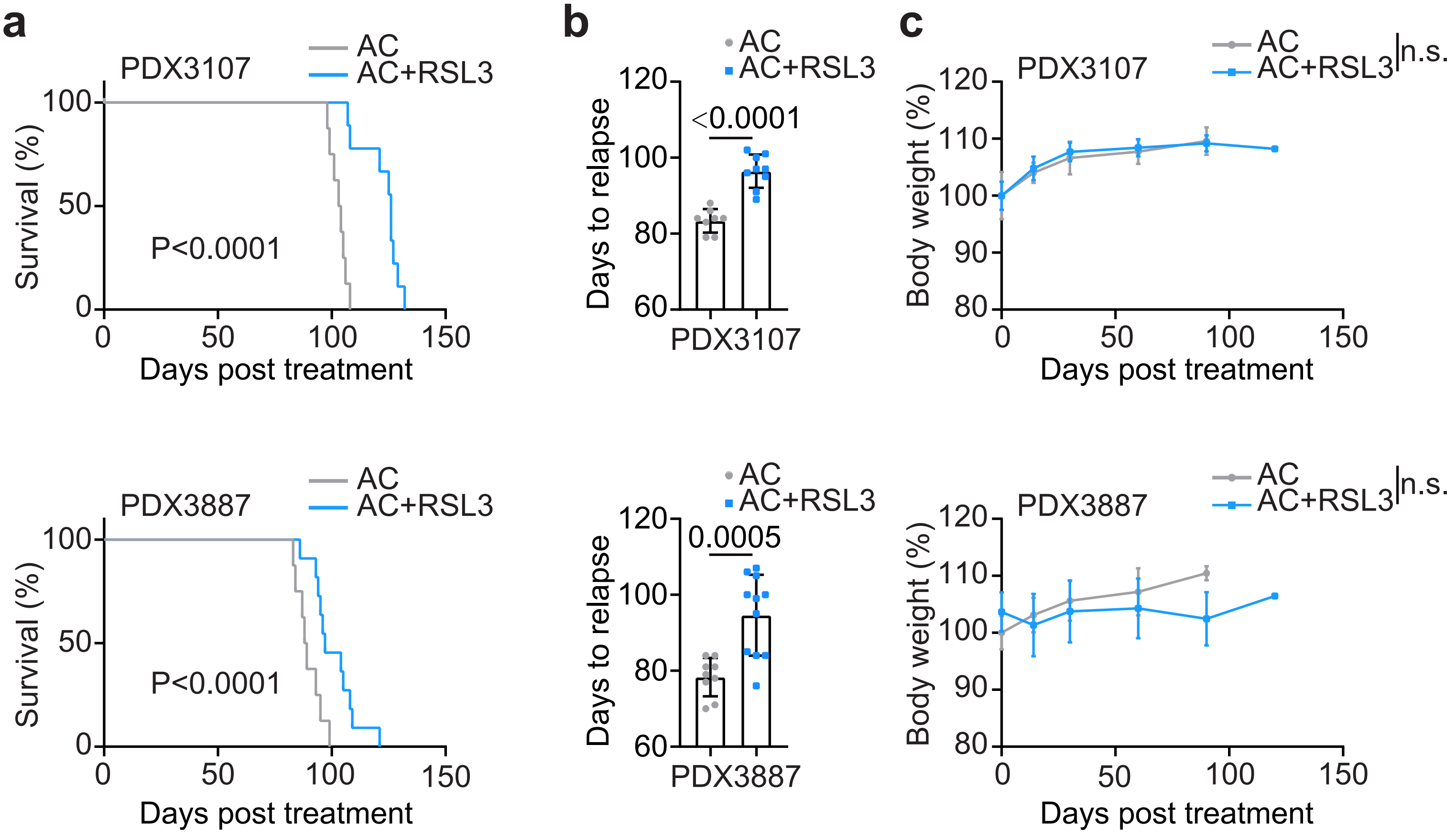

### Supplementary Fig 6

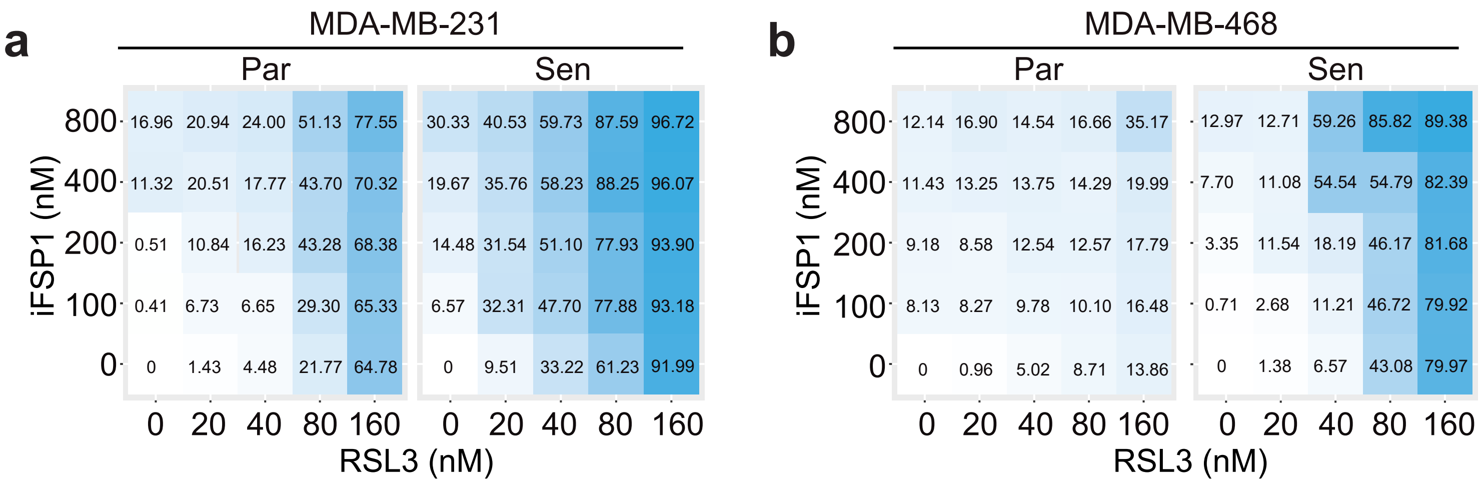

### Supplementary Fig 7

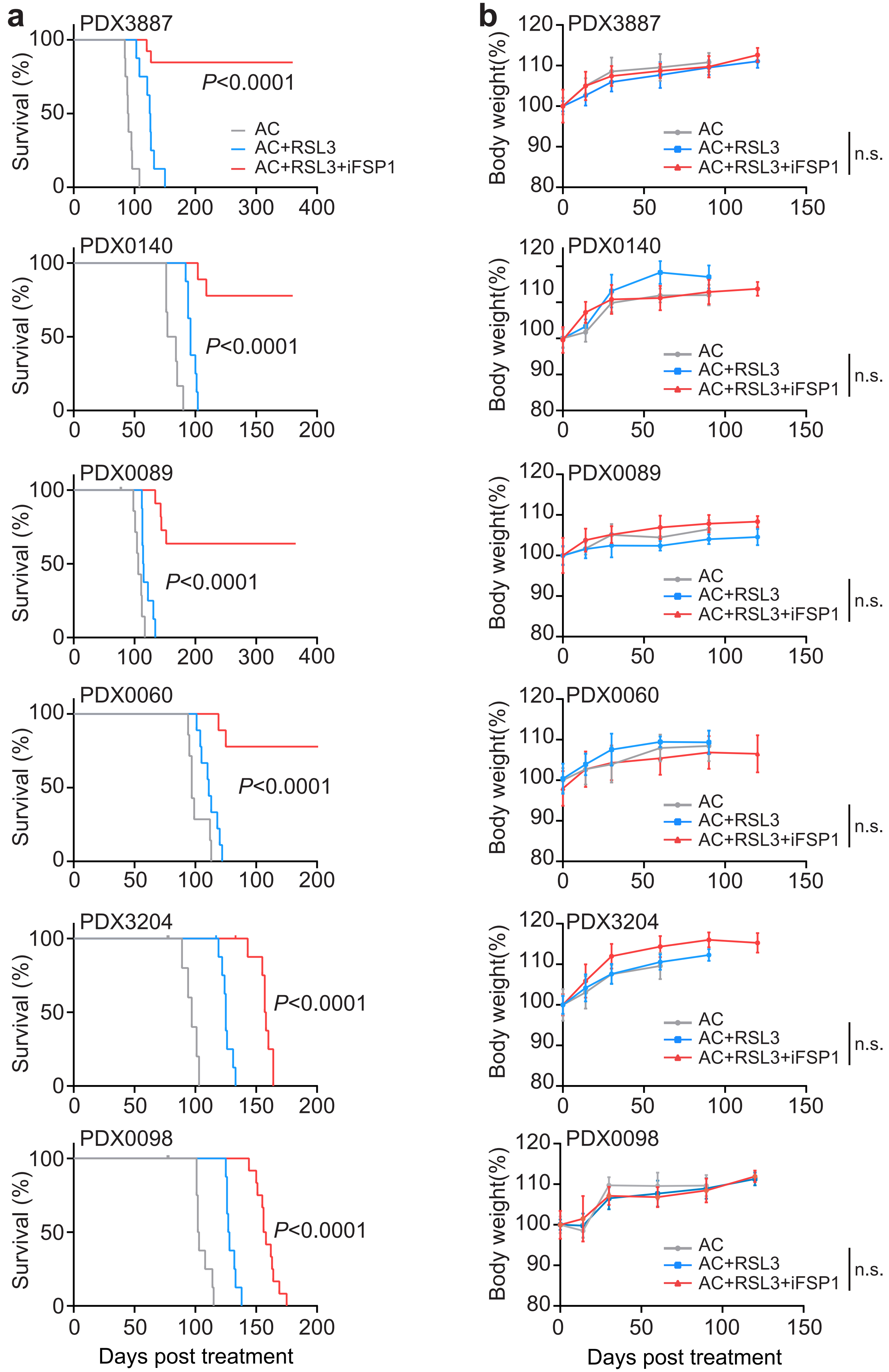

### Supplementary Fig 7
(*continued*)

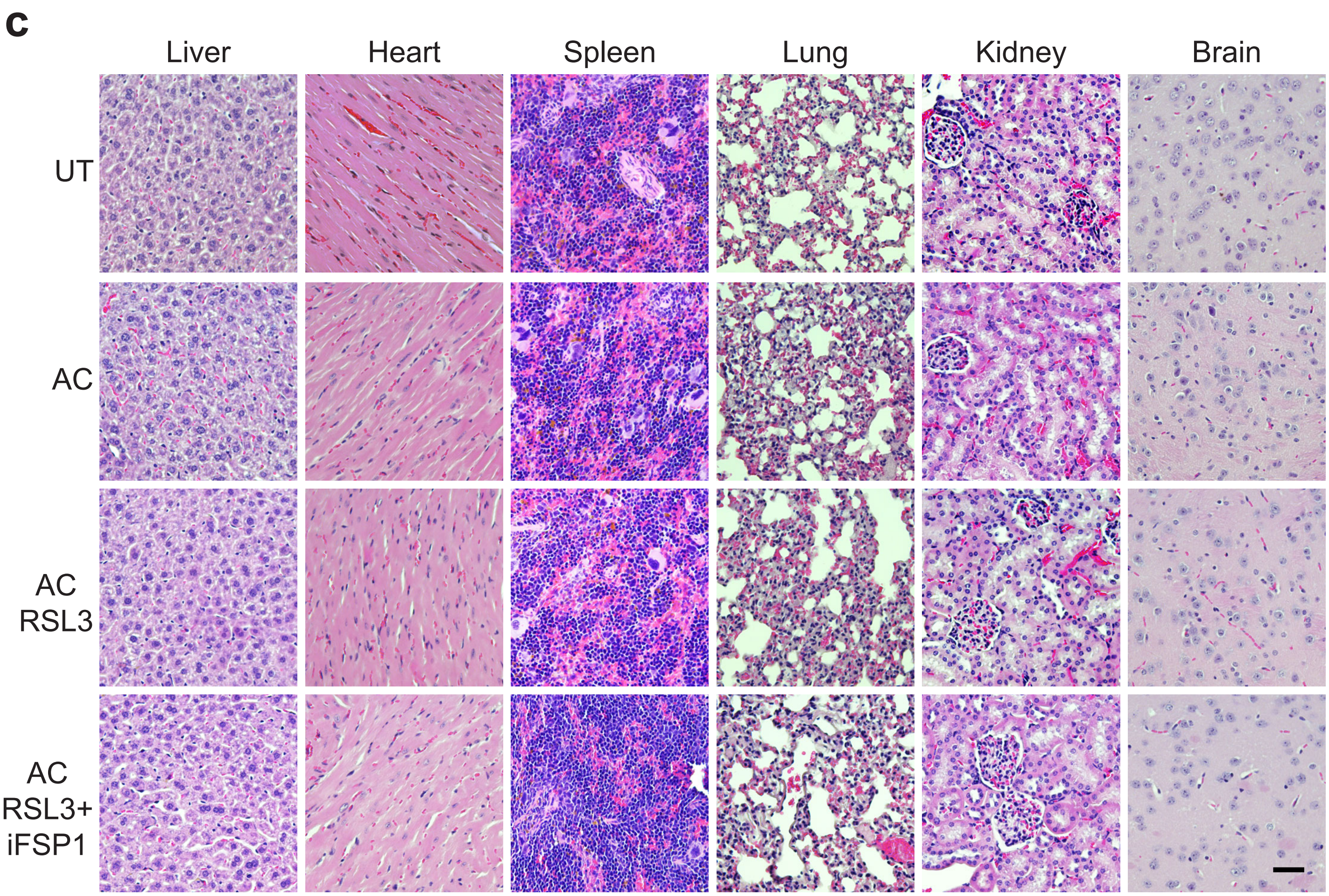

### Supplementary Fig 8

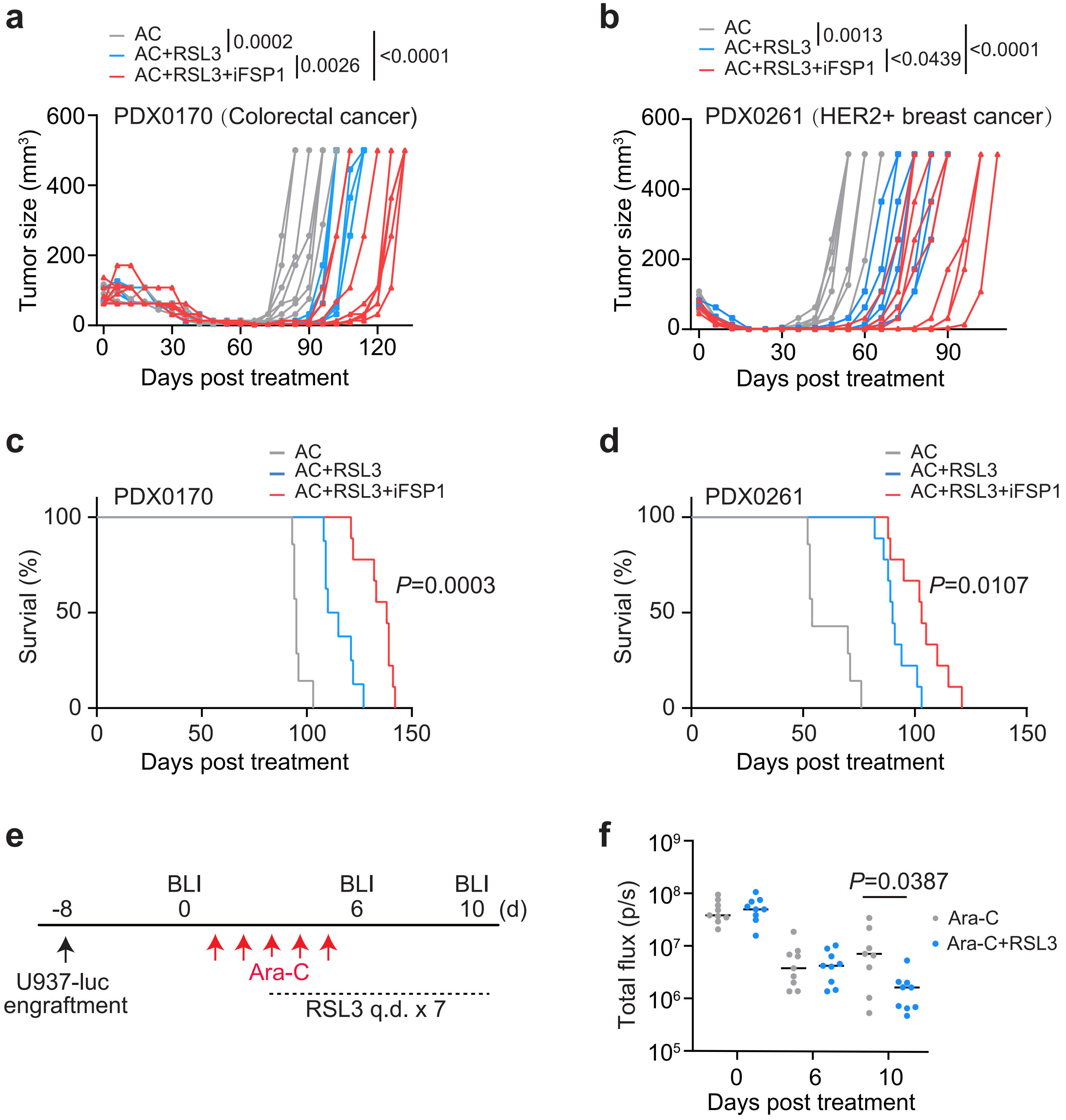

### Supplementary Fig 9

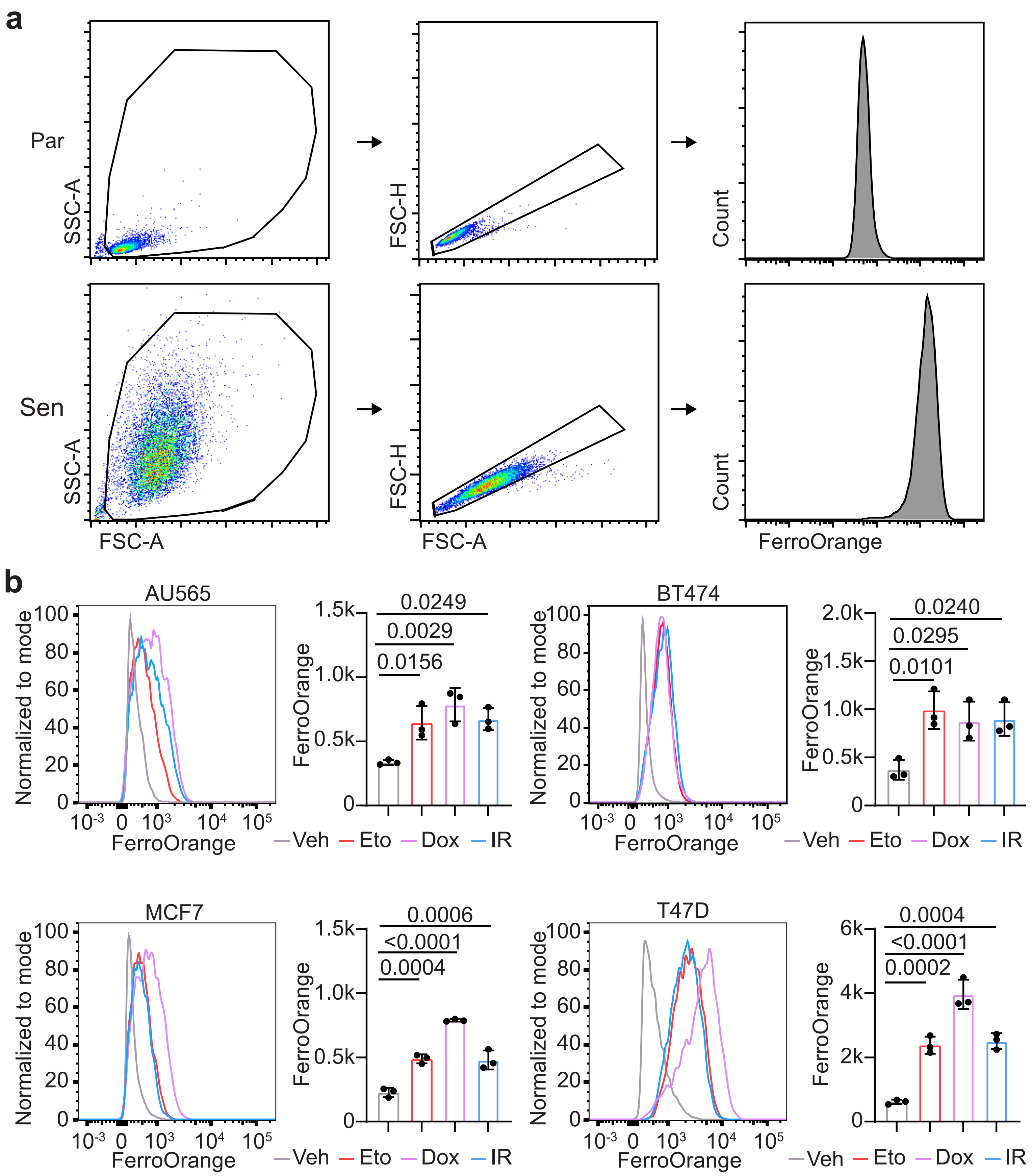

### Supplementary Fig 10

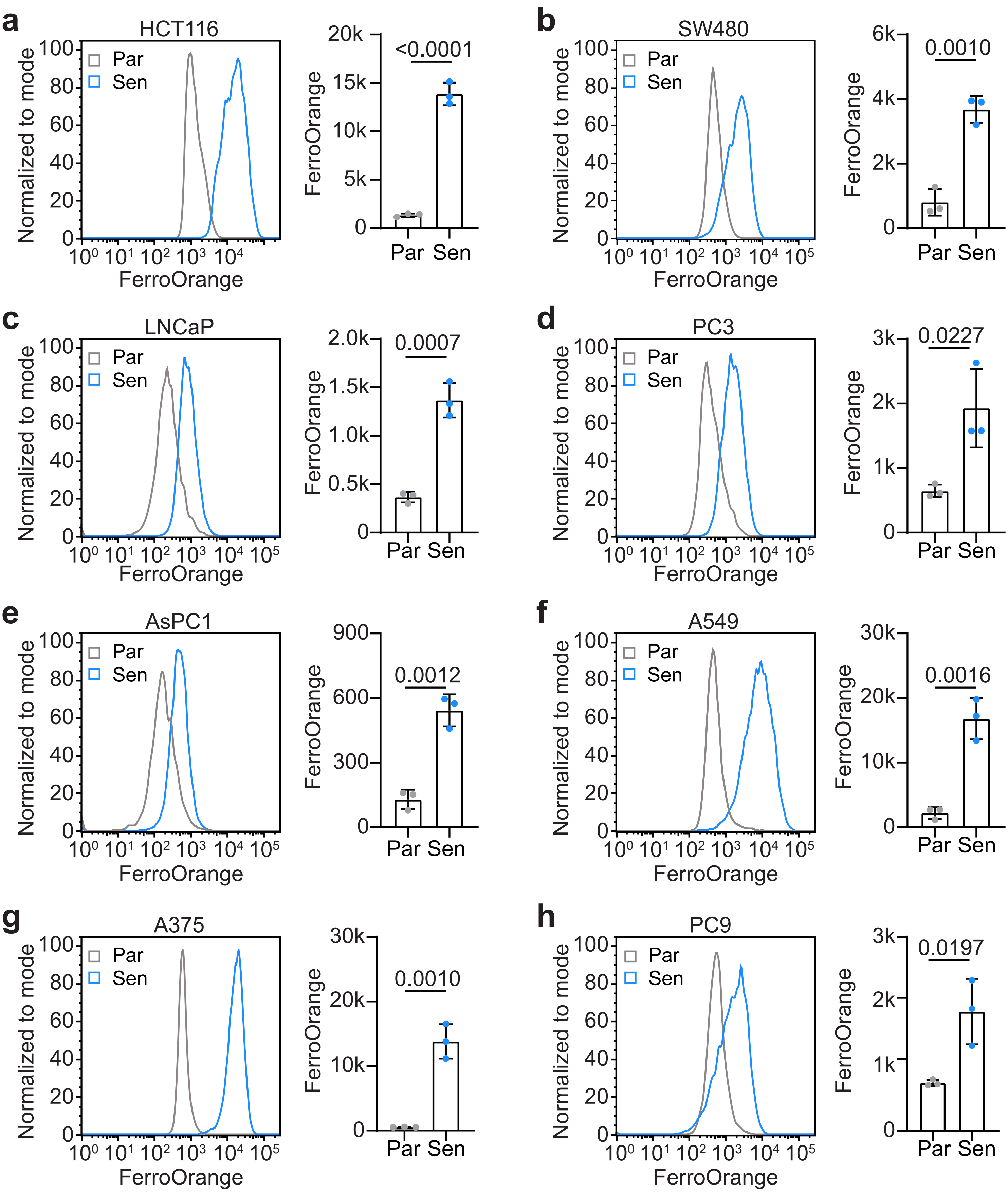

### Supplementary Fig 11

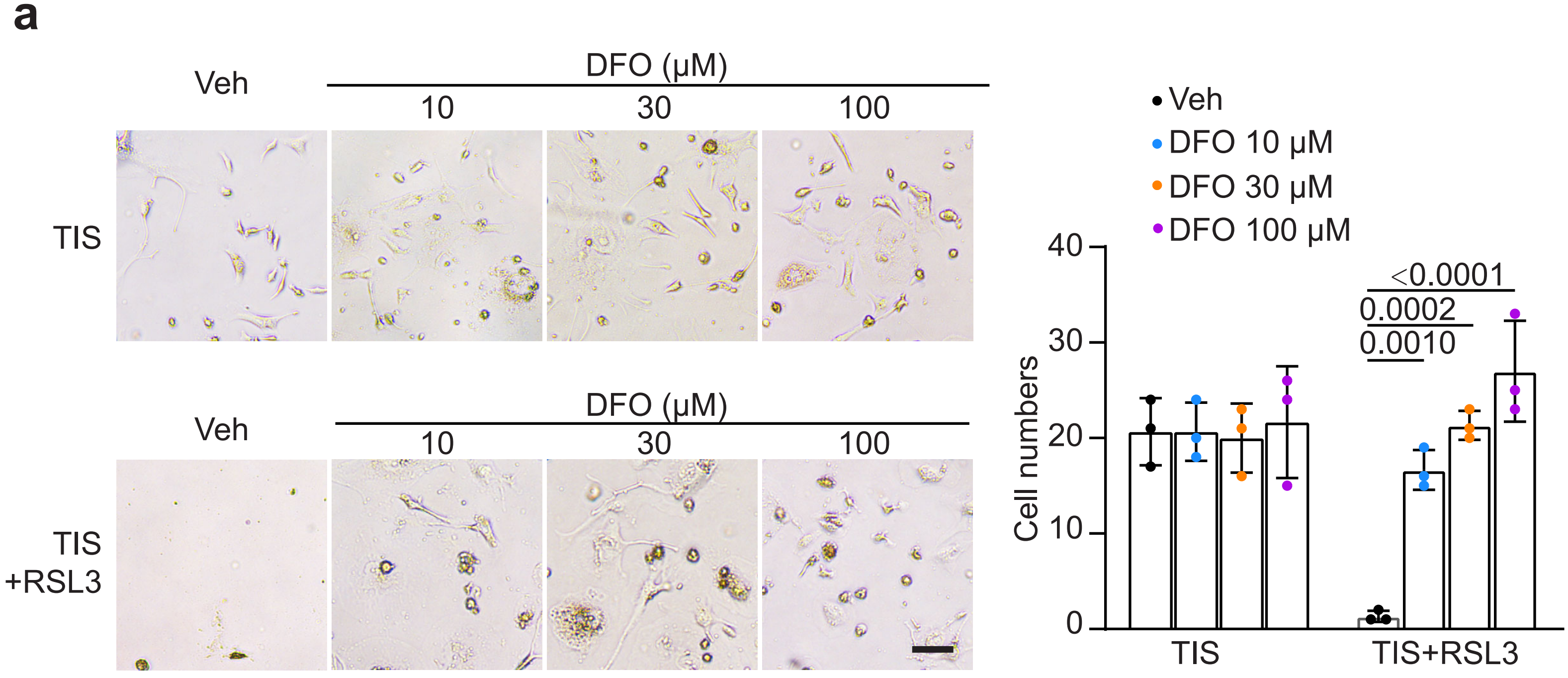


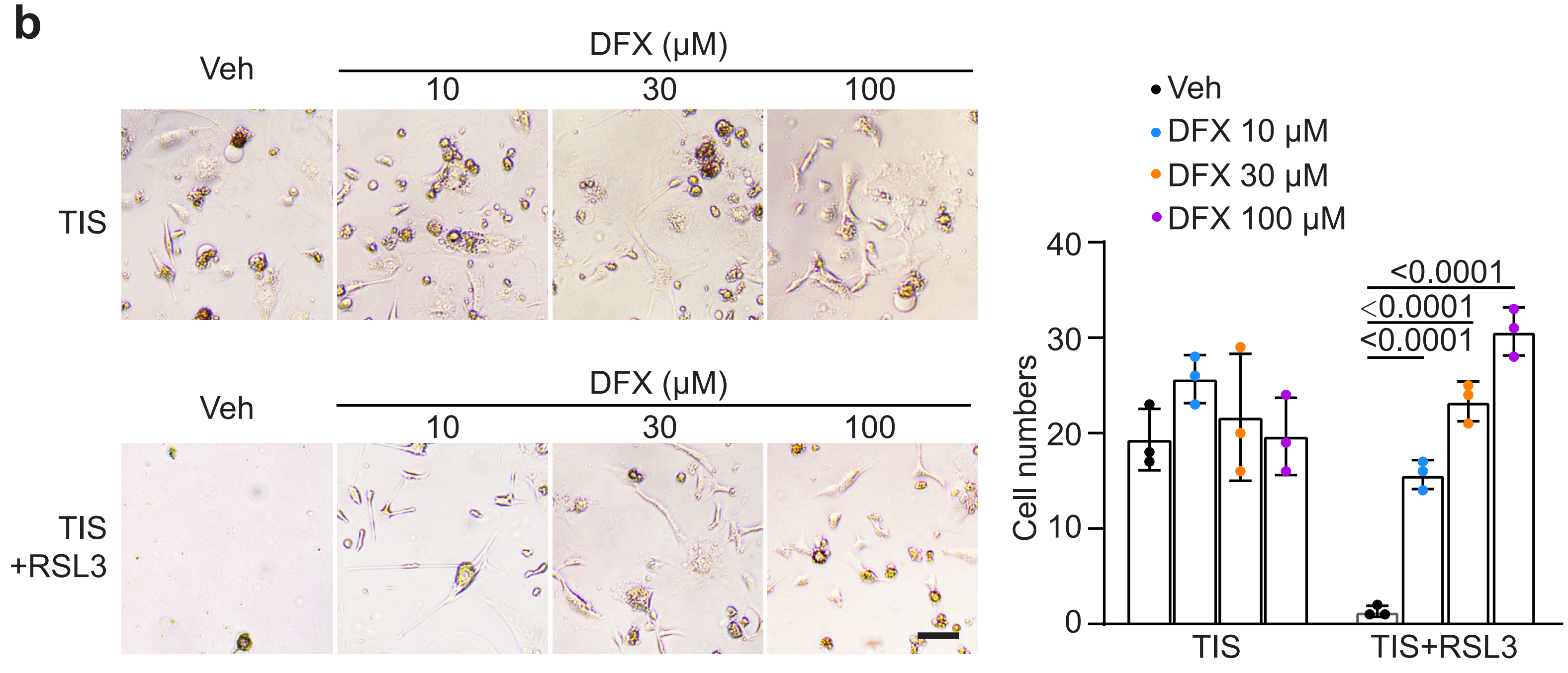

### Supplementary Fig 12

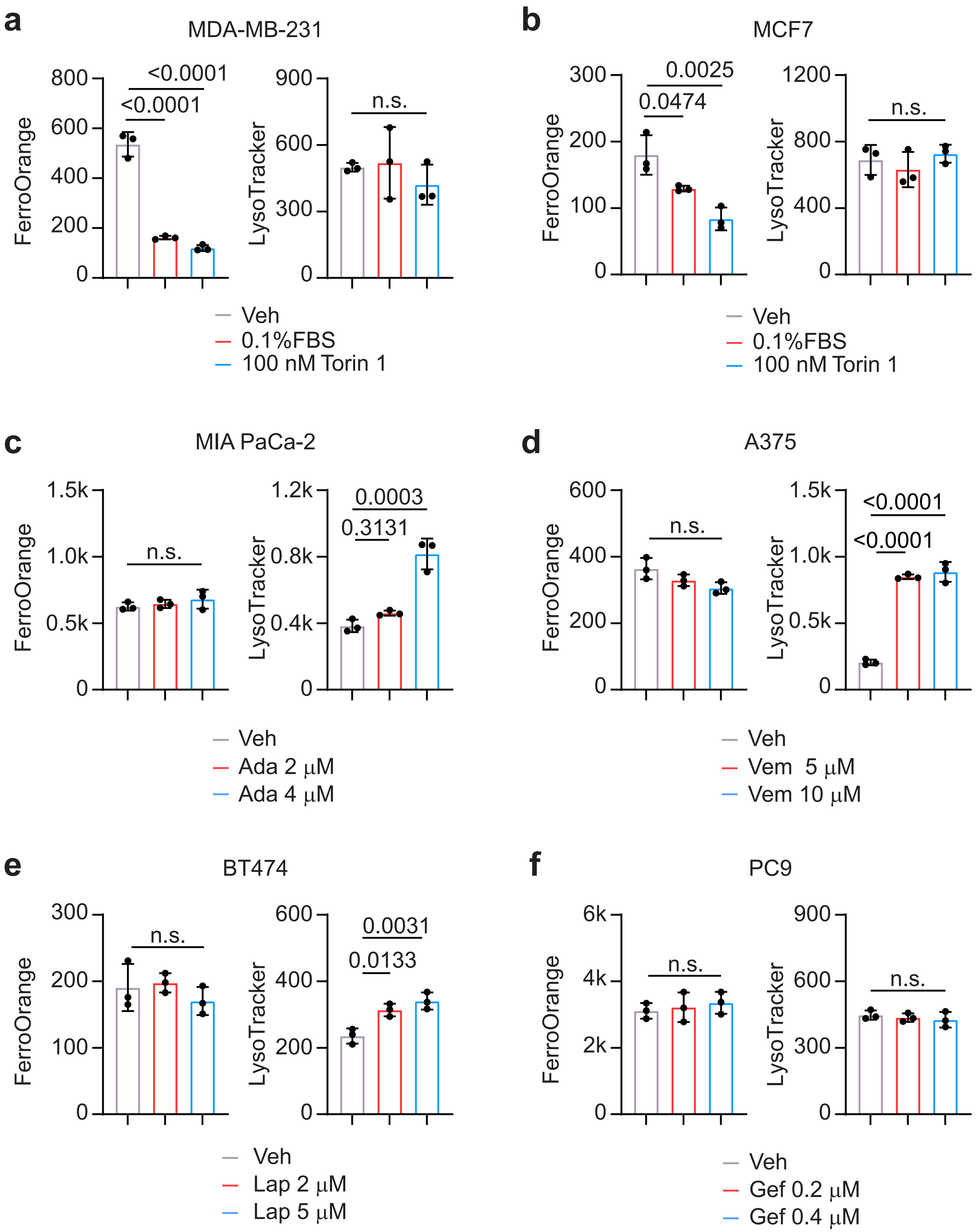

### Supplementary Figure 13

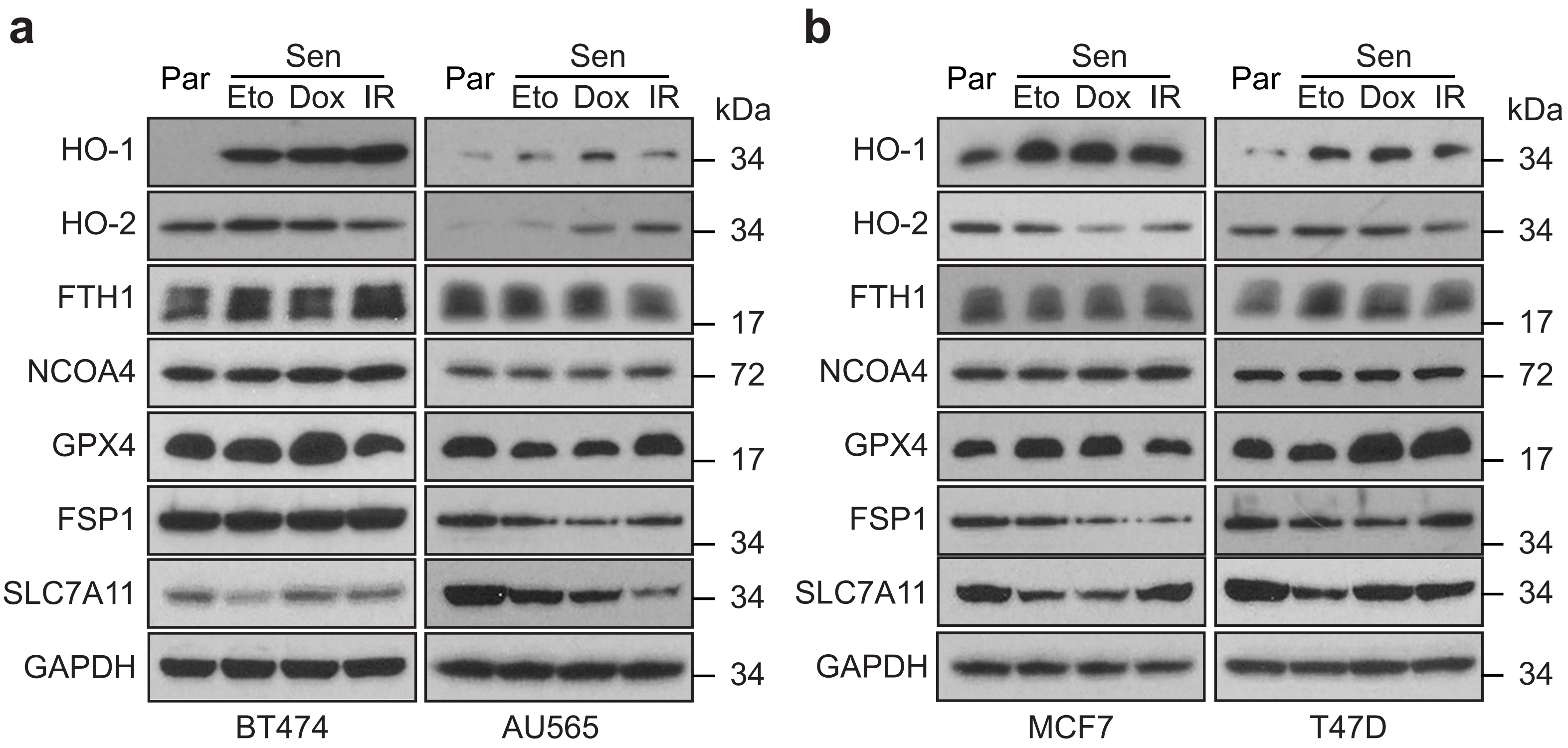
